# Transcriptional control of T cell tissue adaptation and effector function in infants and adults

**DOI:** 10.1101/2025.02.01.636039

**Authors:** Peter A. Szabo, Hanna M. Levitin, Thomas J. Connors, David Chen, Jenny Jin, Puspa Thapa, Rebecca Guyer, Daniel P. Caron, Joshua I. Gray, Rei Matsumoto, Masaru Kubota, Maigan Brusko, Todd M. Brusko, Donna L. Farber, Peter A. Sims

## Abstract

The first years of life are essential for the development of memory T cells, which rapidly populate the body’s diverse tissue sites during infancy. However, the degree to which tissue memory T cell responses in early life reflect those during adulthood is unclear. Here, we use single cell RNA-sequencing of resting and *ex vivo* activated T cells from lymphoid and mucosal tissues of infant (aged 2-9 months) and adult (aged 40-65 years) human organ donors to dissect the transcriptional programming of memory T cells over age. Infant memory T cells demonstrate a unique stem-like transcriptional profile and tissue adaptation program, yet exhibit reduced activation capacity and effector function relative to adults. Using CRISPR-Cas9 knockdown, we define Helios (*IKZF2*) as a critical transcriptional regulator of the infant-specific tissue adaptation program and restricted effector state. Our findings reveal key transcriptional mechanisms that control tissue T cell fate and function in early life.

---

The maturation of adaptive immunity in early life is essential for the establishment of protective immune memory that can last a lifetime. The first years of age represent an intense period of exposure to novel antigens that generate memory T cells required for orchestrating acquired immunity and vaccine-induced protection against infectious disease. However, infants exhibit reduced or diminished responses to ubiquitous pathogens and vaccines relative to adults^1^. While recent studies uncovered distinct activation pathways and effector profiles for infant and adult T cells^2,3^, the underlying mechanisms for this discrepancy in functional capacity remain unknown. Understanding the interplay between the maturation and function of T cells in infancy is necessary for advancing vaccine strategies and immunotherapies targeted to early life.

T cell responses in infancy are distinct from those in adulthood. While initial studies describe infant T cells as intrinsically impaired in effector functions relative to adults, an updated paradigm holds that infant T cells exhibit distinct effector responses that are adapted to the unique demands of early life^4^. Infant naïve T cells preferentially produce T helper type 2 (TH2) cytokines or chemokines (e.g., CXCL8) instead of pro-inflammatory (TH1) cytokines upon activation^5–8^. We and others previously showed that infant naïve T cells are more sensitive to T cell receptor (TCR)-stimulation, exhibit augmented proliferative responses, and demonstrate biased differentiation towards short-lived effector cells compared to adults^9–12^. Features of this infant-specific response may be traced to transcriptional programming or distinct progenitors within the naïve T cell pool, predisposing cells towards mounting effector responses during infections at the expense of forming memory^3,10,11,13^. However, the mechanisms governing effector responses of memory T cell populations that are formed during infancy are not well understood.

We previously showed that the generation of T cell memory in early life begins in tissues, particularly in mucosal sites such as the lungs and intestines that represent the frontlines of antigen exposure, while the vast majority of T cells in the blood remain naïve^14–16^. These infant tissue memory T cells predominantly exhibit an effector memory (TEM) phenotype with markers of tissue residency (e.g., CD69 and CD103) but show decreased expression of tissue homing/adhesion molecules and reduced production of inflammatory mediators upon stimulation compared to older children and adults^16,17^. We recently defined transcriptional profiles of tissue memory T cells during infancy and childhood including expression of transcription factors (TFs) associated with T cell development^16^. How these and other transcriptional regulators control maturation and effector responses of tissue memory T cells during infancy is not known.

Here, we use single cell RNA-sequencing (scRNA-seq) of resting and *ex-vivo* activated T cells from lymphoid and mucosal tissues of infant (aged 2-9 months) and adult (aged 40-65 years) human organ donors to dissect the transcriptional programming of tissue memory T cells in early life. We apply a consensus-implementation of single cell Hierarchical Poisson Factorization (consensus-scHPF)^18^ to define transcriptional states associated with T cell activation, effector function, and tissue adaptation across tissues in infants and adults. We find that relative to adults, infant tissue memory T cells demonstrate a stem-like transcriptional profile (*TCF1*, *LEF1*, *SOX4*), yet exhibit restricted transcriptional responses to TCR-mediated stimulation. We elucidate unique tissue-associated transcriptional states between infant and adult tissue memory T cells and uncover drivers of these programs by gene regulatory network reconstruction. Using CRISPR-Cas9 knockdown in primary tissue T cells, we define Helios (*IKZF2*) as a critical regulator of an infant-specific tissue adaptation program and demonstrate that Helios also restricts infant T cell effector function after stimulation. Together, our results reveal key mechanisms by which age impacts T cell fate and function, with important implications for targeting T cell responses during the formative years of infancy.

## RESULTS

### A single cell transcriptional map of T cell activation across infant and adult tissues

To define the transcriptional programming of T cells across tissues in early life, we performed scRNA-seq on T cells from lymphoid and mucosal sites in infants (2-9 months old) and adults (40-65 years old) (**Supplementary Table 1**). Blood and tissues were obtained from deceased organ donors at the time of life-saving transplantation, including lymphoid organs (bone marrow, spleen, tonsil, intestinal Peyer’s patches, and lung-, jejunum-, and colon-associated lymph nodes) and mucosal/barrier tissues (lungs, jejunum, ileum, colon). Purified T cell populations from these sites were obtained by magnetic selection and cultured overnight in media alone (“resting”) or stimulated with anti-CD3 and anti-CD28 antibodies (“activated”) prior to single cell sequencing using the 10x Genomics Chromium platform (**Fig. 1a**). We merged this dataset with our previous study of resting and activated T cells from adult human organ donors and living blood donor volunteers^19^, for a total of ∼275,000 single cell profiles of T cells across 12 tissues.

**Figure 1:**
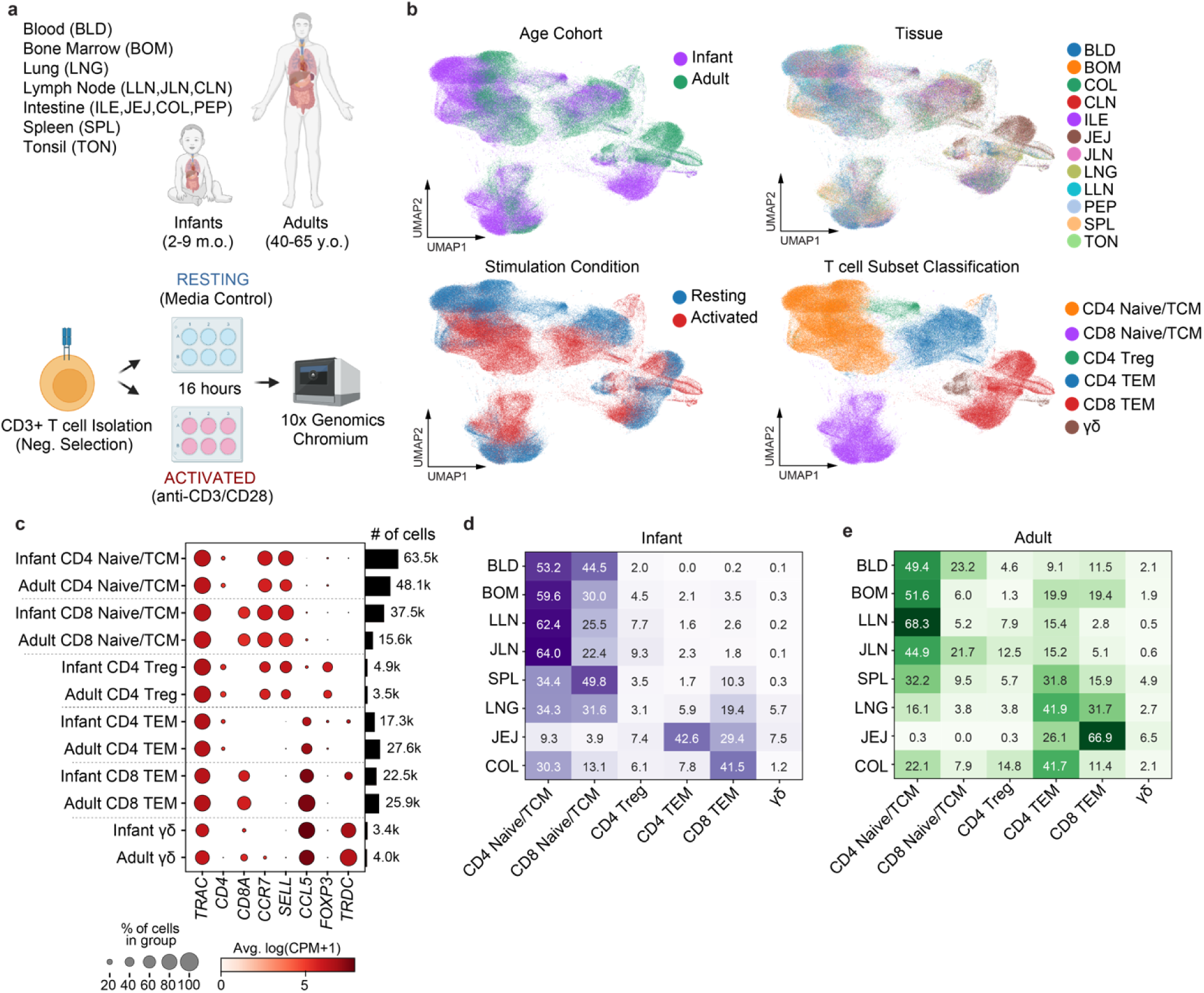
A single cell transcriptomic map of resting and activated T cells from human lymphoid and mucosal tissues. **a**) Schematic of scRNA-seq experimental design and workflow including T cell isolation from infant and adult tissues using negative selection, overnight rest or TCR-mediated stimulation with anti-CD3 and anti-CD28 antibodies, and single cell encapsulation with the 10x Genomics Chromium system. **b**) UMAP embeddings of merged scRNA-seq profiles from resting and activated T cells from all samples, colored by age cohort, tissue of origin, stimulation condition and T cell subset classification. **c**) Dot plot displaying expression of T cell lineage defining markers across T cell subsets for infants and adults. Color intensity reflects mean gene expression by group and dot size reflects percentage of cells in each group expressing indicated marker genes. Number of cells in each T cell subset across all donors and tissues for each age cohort is indicated in the bar plot on the right. **d**) Heatmap indicating percentage of cells for each T cell subset within a tissue (i.e., row-normalized) for infants where color intensity indicates higher frequencies. **e**) Heatmap same as (**d**) but for adults. Abbreviations: blood (BLD), bone marrow (BOM), colon (COL), colon-associated lymph node (CLN), ileum (ILE), jejunum (JEJ), jejunum-associated lymph node (JLN), lung (LNG), lung-associated lymph node (LLN), Peyer’s patch (PEP), spleen (SPL), tonsil (TON).

We first defined T cell subsets for CD4^+^ and CD8^+^ T cells in the merged dataset as naïve/central memory T cells (Naive/TCM), effector memory T cells (TEM), CD4^+^ regulatory T cells (CD4^+^ Tregs), and γδ T cells using a Naïve Bayes classifier (**Supplementary Fig. 1, see Methods**). Visualization of the dataset by uniform manifold approximation and projection (UMAP)^20^ revealed that age cohort and stimulation conditions were dominant sources of transcriptional variability within T cell subsets (**Fig. 1b**). The expression levels of canonical marker genes defining the T cell subsets were highly conserved between infants and adults: CD4^+^ (*CD4)* and CD8^+^ (*CD8A*) naïve/TCM were enriched in lymphoid homing molecules *CCR7* and *SELL* (coding for CD62L); CD4^+^ Tregs uniquely expressed *FOXP3*; CD4^+^ and CD8^+^ TEM highly expressed *CCL5* as a marker of TEM cells defined previously^19^; γδ T cells showed increased expression of *TRDC* and decreased expression of *TRAC*, which encode the eponymous δ-chain or α-chain constant region of the TCR for γδ or conventional αβ T cells, respectively (**Fig. 1c** and **Supplementary Fig. 2**).

For a direct comparison of T cell populations across infant and adult tissues, we focused our analysis on tissue sites that were represented in both age cohorts: blood, bone marrow, lung- and jejunum-associated lymph nodes, spleen, lungs, jejunum and colon. In infants, the vast majority of T cells in blood and lymphoid sites were CD4^+^ and CD8^+^ naïve/TCM, with minor populations of CD4^+^ Tregs, and few TEM or γδ T cells (**Fig. 1d**). Mucosal sites and spleen showed greater proportions of TEM, particularly for CD8^+^ T cells, and the majority of intestinal T cells were either CD4^+^ or CD8^+^ TEM, consistent with our previous findings^14–16^. By contrast, in adults CD4^+^ and CD8^+^ TEM predominated relative to naïve/TCM in mucosal sites and the spleen (**Fig. 1e**). Notably, we observed much lower proportions of CD8^+^ naïve/TCM as compared to CD4^+^ naïve/TCM T cells in infants across most tissues relative to adults (**Fig. 1d,e**). These findings are consistent with an exponential increase in T cell memory observed across infancy and childhood compared to adults^16^.

### Infant TEM exhibit distinct a stem-like transcriptional state relative to adults

We directly investigated changes in gene expression between infant and adult T cells in the resting state using pairwise differential expression analysis across all donors and tissues with adequate representation for each subset (**see Methods**). CD4^+^ and CD8^+^ TEM exhibited a large number of differentially expressed genes (193 and 173, respectively) between the age cohorts (**Extended Data Fig. 1a** and **Supplementary Table 2**). Many of these differentially expressed genes were shared across CD4^+^ and CD8^+^ TEM in infants (**Extended Data Fig. 1b**), demonstrating a conserved transcriptional state in early life. We also detected shared genes expressed by CD4^+^ and CD8^+^ naïve/TCM that were upregulated in adults (**Extended Data Fig. 1c**), which was likely due to increased frequencies in TCM populations^16^, whose profiles could not be readily distinguished from naïve T cells by gene expression alone^21^.

Infant CD4^+^ and CD8^+^ TEM showed significant upregulation of genes encoding TFs linked to T cell stemness/self-renewal and quiescence, including *TCF7* (TCF1)*, LEF1* and *KLF2* (**Fig. 2a,b**)^22–24^. *SOX4*, which cooperates with TCF/LEF family TFs in the Wnt signaling pathway^25^, and *IKZF2* (Helios), typically associated with Treg differentiation and function^26,27^, were also upregulated in infant TEM across sites. Infant CD4^+^ TEM were enriched for expression of the TH2-driving TF *GATA3*^28^, while infant CD8^+^ TEM expressed high levels of the resident- and effector-associated TFs *ZNF683* (Hobit) and *ID3*^29,30^ relative to adults. We also observed an increase in transcripts associated with innate-like T cells (*ZBTB16*, *NCR3*, *KLRB1*, *FCER1G*)^31^ in infants across tissues (**Fig. 2a,b**). Lastly, we found increased expression of genes encoding T cell co-stimulatory or inhibitory surface molecules (*CD27, CD28, CD38, KLRG1*) in both lineages of infant TEM relative to those in adults.

**Figure 2:**
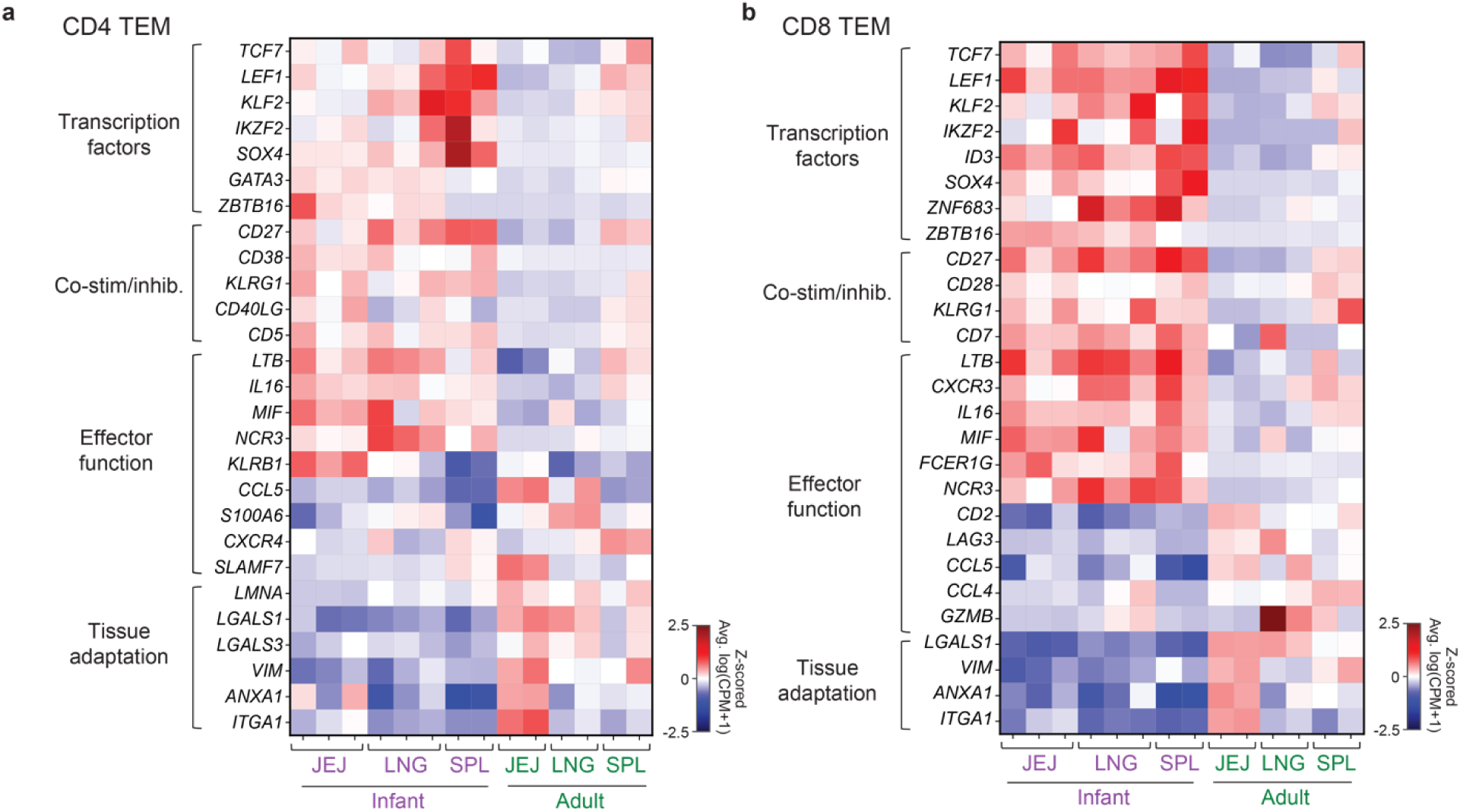
Differentially expressed genes across infant and adult TEM across tissues. **a**) Heatmap of selected differentially expressed genes in infant versus adult CD4^+^ TEM from indicated tissues in the resting condition. Color intensity reflects Z-scored average log(counts per million+1) expression by row. **b**) Heatmap of differentially expressed genes in infants and adults as in (**a**) but for CD8^+^ TEM.

Expression of genes associated with T cell effector function were variably expressed between infant and adult TEM (**Fig. 2a,b**). Across tissues, genes coding for cytokines (*LTB, MIF, IL16*) and chemokine receptors (*CXCR3, CXCR4*) were upregulated in infant TEM, while chemokines (*CCL4*, *CCL5*) and mediators of cytotoxicity (*GZMB, SLAMF7*) were upregulated in adults. Infant TEM also exhibited upregulated expression of genes associated with activation (*CD38*, *CD40LG*) and effector T cell fate (*KLRG1*), consistent with ongoing activation and effector differentiation in infants encountering many new antigens. By contrast, adult CD4^+^ and CD8^+^ TEM showed increased expression of transcripts associated with tissue adaptation and adhesion, including *LGALS1* (galectin-1), *ANXA1* (annexin-1), *VIM* (vimentin), and *ITGA1* (CD49a)^19,32^.

Given the critical and multi-faceted roles of TCF1 and LEF1 in T cell identity and function^33^, we sought to validate the augmented expression of both TFs in infant versus adult TEM on the protein level by flow cytometry. The expression of TCF1 and LEF1 by CD8^+^ TEM was significantly increased in infants compared to adults in the spleen, while only LEF1 was increased on CD4^+^ TEM (**Extended Data Fig. 1d,e**). Taken together, our findings demonstrate that infant CD4^+^ and CD8^+^ TEM exhibit a distinct transcriptional state with increased expression of transcriptional regulators of stemness and memory differentiation relative to adults.

### Consensus-scHPF reveals unique signatures of tissue adaptation and effector function across infant and adult tissue T cells

To uncover unique gene expression programs between infant and adult tissue T cells, we utilized consensus-scHPF^18^, a probabilistic Bayesian factorization method for the *de novo* discovery of latent transcriptional co-expression signatures or “factors” in scRNA-seq data (diagrammed in **Fig. 3a**). We applied consensus-scHPF to all infant and adult tissue T cells and identified discrete factors defined by the top genes in the consensus gene score matrix (**Fig. 3b,c** and **Supplementary Table 3**). To assess whether a given factor was associated with specific features of dataset, we performed multivariate linear regression using age cohort (infant or adult), tissue localization (lymphoid or mucosal), T cell subset (naïve/TCM or effector), T cell lineage (CD4 or CD8), and activation condition (resting or activated) as covariates and plotted regression coefficients for each comparison (**Fig. 3d**). In total, we identified 18 distinct signatures corresponding to T cell subsets, metabolism, tissue adaptation, activation states, and/or effector functions across infant and adult tissue T cells.

**Figure 3:**
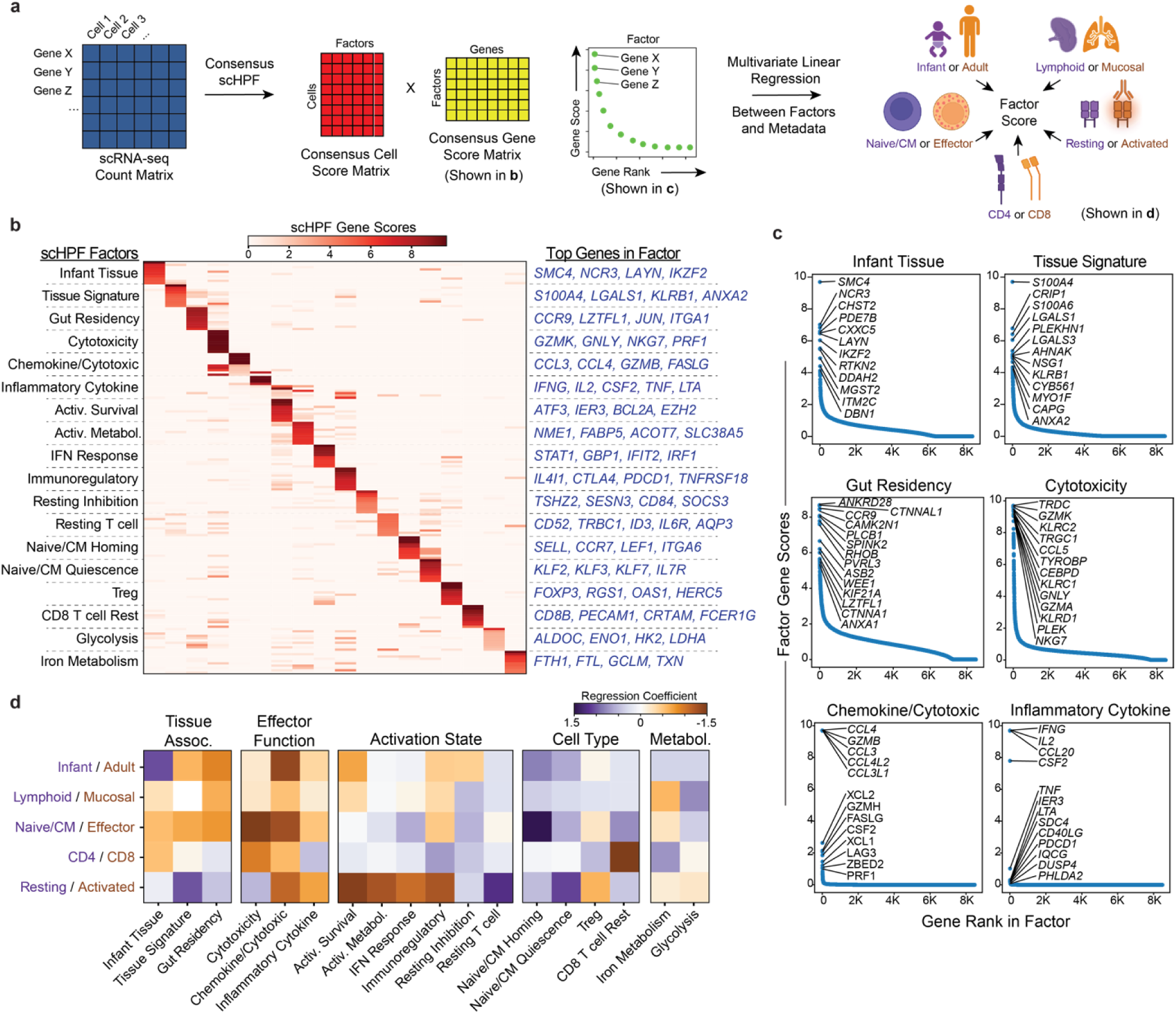
Consensus single cell Hierarchical Poisson Factorization (scHPF) reveals transcriptional co-expression patterns in infant and adult T cells. **a**) Schematic of consensus-scHPF analysis to identify co-expression patterns or “factors” across the dataset and multivariate linear regression to associate factors with metadata. **b**) Heatmap showing scHPF gene scores of the top 10 genes in each factor. Selected genes for each factor are indicated to the right and a ranked list of the top 100 genes in all factors is provided in **Supplementary Table 3**. **c**) Dot plots showing the rank and gene score for genes in selected scHPF factors with labels for the top genes. **d**) Heatmap of regression coefficients for multivariate linear regression between scHPF factor cell scores and covariates: age cohort (infant/adult), tissue localization (lymphoid/mucosal), T cell subset (naïveCM/effector), T cell lineage (CD4/CD8), or activation conditions (resting/activated).

Consensus-scHPF revealed three factors associated with tissue localization and adaptation that differed between infants and adults. The first tissue factor (“Infant Tissue”) was defined by transcripts that were highly differentially expressed in infant TEM relative to adults from our previous analysis (from **Fig. 2**), including *SMC4, NCR3, CXXC5, LAYN,* and *IKZF2,* and was strongly enriched in infant CD8^+^ TEM from mucosal tissues (**Fig. 3c,d**). The second tissue factor (“Tissue Signature”) was characterized by genes that we previously identified as a signature of T cells residing in tissues compared to the blood^19,32^, including *S100A4/6*, *CRIP1*, *LGALS1*, *KLRB1*, and *ANXA2,* and was associated with adult TEM (**Fig. 3c,d**). The third tissue factor (“Gut Residency”) was strongly biased towards adult TEM and was distinguished by markers of intestinal homing and adhesion (*CCR9*, *ITGA1, CTNNA1*)^32,34^ and tissue-resident memory T cell (TRM) development (*AHR, JUN, FOSB*)^35,36^ (**Fig. 3c,d**).

Three scHPF factors were associated with distinct T cell effector states. The “Cytotoxicity” factor was defined by cytolytic molecules *GZMK, GNLY, GZMA, NKG7* and *PRF1*, and was highly enriched in mucosal CD8^+^ TEM associated with the resting condition (**Fig. 3c,d**), reflecting a poised cytotoxic state. Relatedly, the “Chemokine/Cytotoxic” factor included highly ranked transcripts for cytotoxic mediators (*GZMB*, *GZMH, FASLG, PRF1*) and potent chemoattractants (*CCL3*, *CCL4, CCL3L1, CCL3L3)*, and was strongly associated with mucosal CD8^+^ TEM in the activated condition (**Fig. 3c,d**). By contrast, the “Inflammatory Cytokine” factor was biased towards CD4^+^ TEM and characterized by genes for *IFNG, IL2, TNF*, *CSF2*, and *LTA* (**Fig. 3c,d**). Importantly, all three factors relating to effector functions and indeed most factors associated with activated conditions in general (e.g., Activation/Survival, Immunoregulatory, Treg), were more strongly associated with adults compared to infants (**Fig. 3d**).

### Infant T cells demonstrate a reduced activation capacity relative to adults

We uncovered several scHPF factors related to T cell activation and effector function that were more strongly associated with adult T cells compared to those in infancy. To interrogate these apparent differences in functional capacity, we investigated the expression of the top genes in both Inflammatory Cytokine and Chemokine/Cytotoxic factors across TCR-simulated CD4^+^ and CD8^+^ TEM from all paired tissues in infants and adults. We found moderate differences in the expression of genes in the Inflammatory Cytokine factor in CD4^+^ TEM between age groups, but strikingly increased expression of these genes among CD8^+^ TEM in adults (**Fig. 4a**). For the Chemokine/Cytotoxic factor, we also observed a prominent increase in expression for its top genes in adult T cells compared to infants, especially for CD8^+^ TEM (**Fig. 4b**). As orthogonal confirmation, we directly assessed the functional capacity of infant and adult TEM via intracellular cytokine staining by flow cytometry after a short term *ex-vivo* stimulation. Both splenic CD4^+^ and CD8^+^ TEM in adults exhibited much greater frequencies of IFNγ, IL-2 and TNFα-producing cells relative to infants (**Fig. 4c,d**), consistent with our findings from scRNA-seq. We also found increased intracellular production of the cytotoxic mediator granzyme B from unstimulated conditions in adult TEM relative to infants, reflecting their augmented poised cytotoxic state suggested by our scHPF analysis (**Fig. 4c,d**).

**Figure 4:**
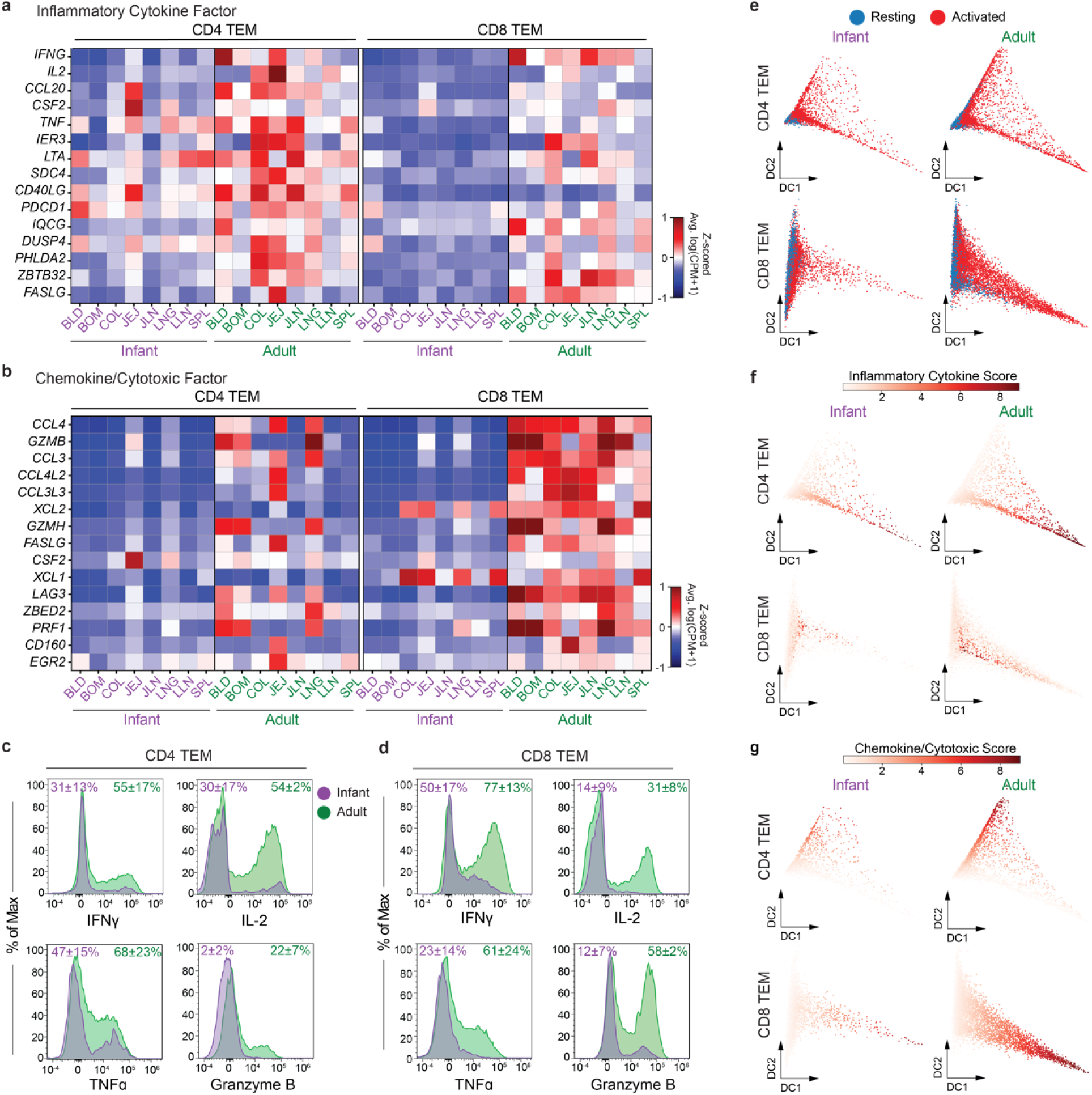
T cell activation capacity and effector function and of CD4^+^ and CD8^+^ TEM in infants and adults. **a**) Heatmap showing z-scored average gene expression as log(counts per million+1) of the top 15 genes in the Inflammatory Cytokine factor in CD4^+^ and CD8^+^ TEM across tissues in infants and adults from activated conditions. **b**) Heatmap as in (**a**) but showing the top 15 genes in the Chemokine/Cytotoxic factor. **c**) Representative histograms of effector molecule expression in infant and adult CD4^+^ TEM from the spleen assessed by intracellular flow cytometry staining. Cytokine (IFNγ, IL-2 and TNFα) expression was evaluated after 4-hour stimulation with PMA and Ionomycin, while granzyme B expression was evaluated in unstimulated conditions. Values represent mean +/- standard deviation of percent positive cells of each target across 3 individual donors in each age cohort. **d**) Representative histograms and percent expression values as in (**c**) but for CD8^+^ TEM. **e**) Diffusion maps of CD4^+^ and CD8^+^ TEM from mucosal tissues (jejunum and lung), with cells colored by activation condition as resting (blue) or activated (red). **f, g**) Diffusion maps as in (**e**) but colored by cell scores for the Inflammatory Cytokine factor (**f**) or Chemokine Cytotoxic factor (**g**) for CD4^+^ TEM (*top*) or CD8^+^ TEM (*bottom*).

For unbiased comparison of T cell activation across age groups, we modeled activation trajectories of T cells from both resting and activated conditions using diffusion maps, for CD4^+^ or CD8^+^ TEM separately. These diffusion maps separated resting T cells (in blue) on the *left* and activated T cells (in red) projecting out to the *right* (**Fig. 4e**). The trajectories for CD4^+^ TEM showed moderate differences in activation between age cohorts, with adults exhibiting an increased number of cells along the activation axis (i.e., x-axis) relative to infants; however, the CD8^+^ TEM trajectories showed strikingly more adult cells along the activation axis compared to infants. Visualizing cell scores of the Inflammatory Cytokine (**Fig. 4f**) and Chemokine/Cytotoxic (**Fig. 4g**) factors from scHPF further highlighted the increased expression of effector signatures on adult CD4^+^ and CD8^+^ T cells compared to infants. These findings collectively demonstrate reduced transcriptional responses to stimulation and a restricted capacity for effector function in infant memory T cells relative to adults.

### Gene regulatory network inference uncovers distinct transcriptional regulators of tissue adaptation in infant and adult tissue T cells

Consensus-scHPF identified three factors related to tissue residency and adaptation that differed across infants and adult mucosal T cells (**Fig. 3c**). The top genes from the Infant Tissue factor were highly enriched among infant CD4^+^ and CD8^+^ TEM (**Extended Data Fig. 2a**), while genes from the Tissue Signature factor were expressed across both infant and adult tissues (**Extended Data Fig. 2b**). By contrast, top genes from the Gut Residency factor were exclusively enriched in the intestinal sites (jejunum, colon, jejunum-associated lymph nodes) of adult CD4^+^ and CD8^+^ TEM (**Extended Data Fig. 2c**), demonstrating a unique transcriptional state.

To investigate the differences in tissue adaptation between age cohorts, we used diffusion maps to model maturation trajectories of resting infant and adult TEM from the intestines (jejunum), where virtually all TEM are tissue-resident in both age groups^16^. For both CD4^+^ and CD8^+^ TEM, the trajectories reflected a continuous transition from an infant state (*left*, purple) to an adult state (*right*, green) and visualizing cell scores for the three tissue-associated scHPF factors on the trajectories highlighted the features of this transition (**Fig. 5a**). The Infant Tissue factor was largely specific to the *left*-most population of infant TEM, whereas high cell scores for the Tissue Signature factor appeared at an intermediate position expressed by subpopulations of both infant and adult TEM. High cell scores for the Gut Residency factor were enriched in adult TEM in the *right*-most population. This analysis demonstrated a continuous tissue adaptation process in infant and adult mucosal TEM, including a shared intermediate state.

**Figure 5:**
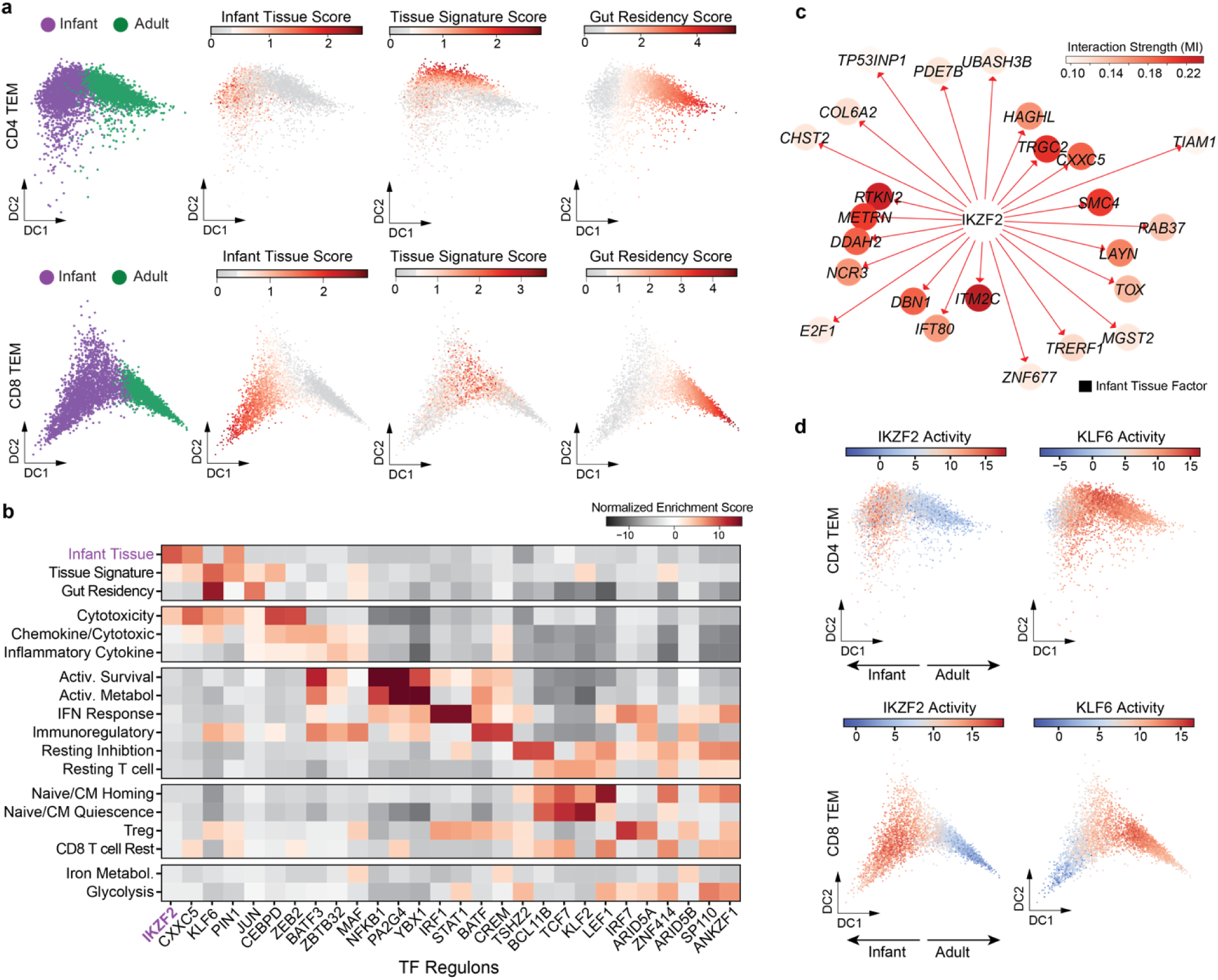
Gene regulatory network reconstruction to identify transcription factors driving tissue adaptation in infant and adult tissue T cells. **a**) Diffusion maps of CD4^+^ and CD8^+^ TEM from the jejunum with cells colored by age cohort as infant (*purple*) and adult (*green*) or by cell scores for the indicated scHPF factors. **b**) Heatmap showing normalized enrichment score from GSEA between the genes in each TF’s regulon and the ranked list of genes for each scHPF factor. The top two TFs with the highest scores for each factor (excluding duplicates) are shown. **c**) TF network of Helios (*IKZF2*) showing inferred TF targets overlapping with the top 50 genes in the Infant Tissue factor. Node color intensity and inverse distance in the network represents interaction strength (mutual information) between TF and target gene. **d**) Diffusion maps from (**a**) but colored by Helios (*IKZF2*) or KLF6 activity determined by VIPER (**see Methods**).

We next sought to identify the putative TFs responsible for driving the functional and tissue-associated transcriptional states in infant and adult T cells. We applied the Algorithm for the Reconstruction of Accurate Cellular Networks (ARACNe), which reverse engineers a gene regulatory network from gene expression data by inferring direct relationships between TFs and their target genes^37,38^. ARACNe generated a set of target genes for each TF, known as a “regulon” (**Supplementary Table 4**). We performed gene set enrichment analysis^39^ between the positively regulated genes in each TF’s regulon and the ranked list of genes for each scHPF factor to associate individual TFs with the cell states defined by each factor. The top two TF regulons with the highest normalized enrichment scores for each scHPF factor are shown in a heatmap in **Fig. 5b**. This analysis identified many TFs previously known to be linked to their respective T cell states including, IRF1 and STAT1 for responses to IFN signaling^40,41^, NFKB1 for T cell activation^42^, IRF7 regulating IFN responses and Tregs^43,44^, ZEB2 for cytotoxic T cell function^45^, and KLF2, TCF1 (*TCF7*) and LEF1 regulating naïve T cell stemness and quiescence^24^.

Importantly, this analysis identified Helios (*IKZF2*) as the top regulator of the Infant Tissue factor and KLF6 as the top regulator of both the Tissue Signature and Gut Residency factors (**Fig. 5b**). We visualized the relationship between the top genes in the tissue adaptation factors with each TF’s regulon using a network, with the mutual information between each TF-gene pair as a measure of their interaction strength. Among the top 50 genes in the Infant Tissue factor, 24 genes were inferred to be regulated by Helios (**Fig. 5c**). Of the top 50 genes in the Tissue Signature and Gut Residency factors, KLF6 was inferred to regulate 21 and 23 genes, respectively (**Extended Data Fig. 2d**).

To assess differences in the activities of TFs across infants and adult T cells, we next performed Virtual Inference of Protein-activity by Enriched Regulon (VIPER) analysis, which utilizes the relative expression of a TF’s up- and down-regulated targets to infer its activity in a given cell^46^. We plotted the TF activities of both Helios and KLF6 on the tissue adaptation trajectories we generated previously and found that Helios activity was restricted to infants, while KLF6 activity was aligned with the transition from infant to adults and increased on all adult CD4^+^ and CD8^+^ TEM (**Fig. 5d**). We next compared the differences in gene expression and activities for each TF in infants versus adults to identify those that were both highly differentially expressed and highly active in each age cohort. Helios was among the top differentially expressed and differentially active TFs in infants CD4^+^ and CD8^+^ TEM (**Extended Data Fig. 2e,f**), along with the other stem-like TFs (TCF1, LEF1, SOX4) that we identified by differential gene expression alone. KLF6 was only mildly enriched in activity and expression in adults, suggesting that differences in the function of KLF6 in infants and adults is not controlled on the level of transcription. Together, these analyses facilitated the discovery of individual TFs associated with T cell states defined by scHPF and identified Helios as putative regulator of the infant-specific T cell tissue adaptation program.

### Helios (*IKZF2*) drives an infant-specific transcriptional program and restrains T cell effector function in early life

To interrogate the functional role of Helios (*IKZF2*) in infant T cells, we first confirmed its expression on the protein level by intracellular staining via flow cytometry. Helios was highly expressed by all subsets of CD8^+^ T cells in infants relative to adults, while only naïve T cells were significantly enriched for Helios in infants among CD4^+^ T cells (**Fig. 6a**).

**Figure 6:**
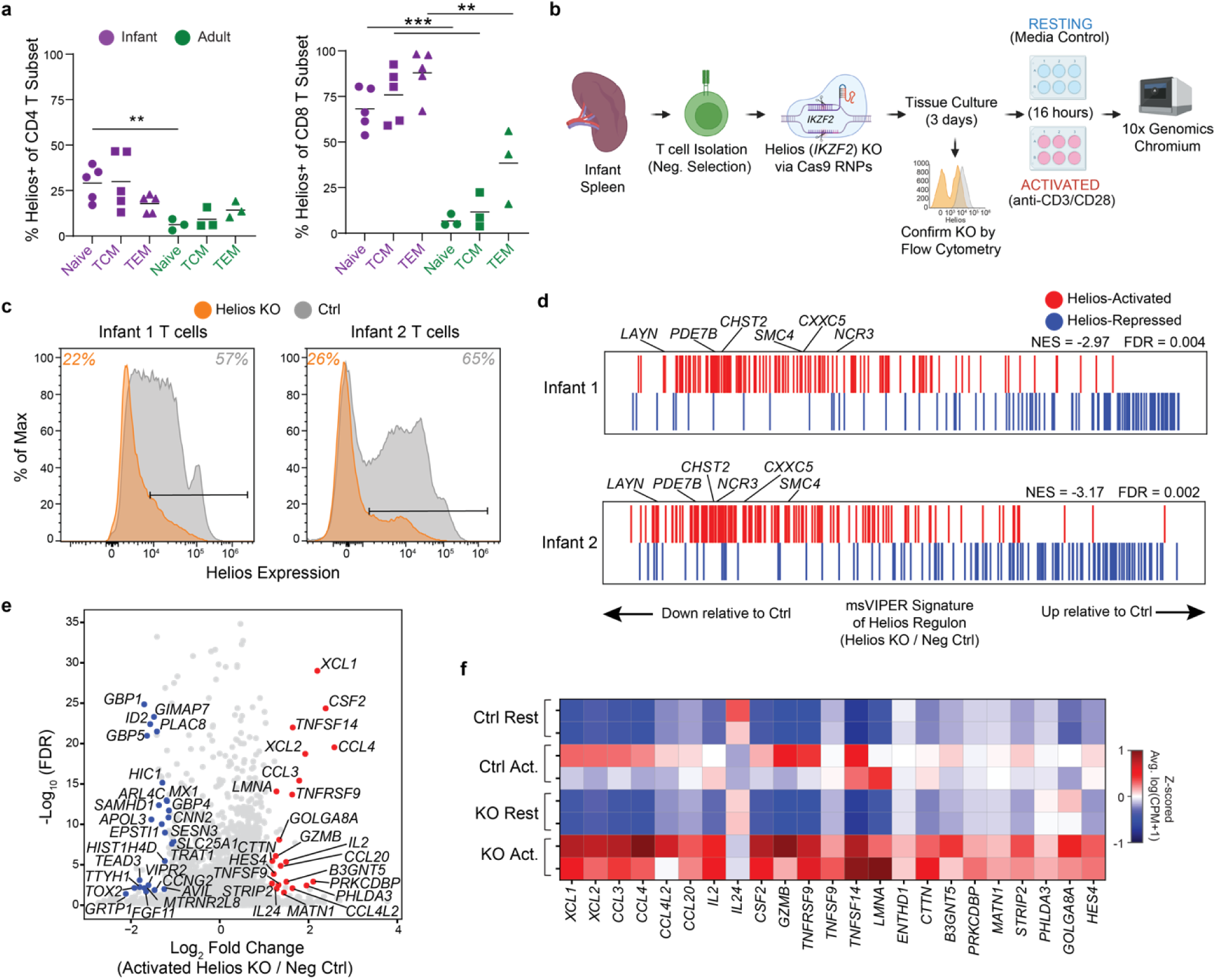
Helios expression and function in infant T cells. **a**) Percentage of Helios-expressing cells among conventional (CD3^+^ γδTCR^-^ FOXP3^-^) CD4^+^ and CD8^+^ T cell subsets from the spleen in infant (n = 5) and adults (n = 3) by flow cytometry. Statistical comparisons between infant and adult subsets made by Students’ *t*-test, where ** denotes *p* < 0.01, *** denotes *p* < 0.001. **b**) Schematic of CRISPR-Cas9 knockout (KO) of Helios (*IKZF2*) in infant splenic T cells after 3 days in culture, overnight rest or activation with anti-CD3 and anti-CD28 antibodies, and single cell sequencing. **c**) Histograms showing protein expression of Helios among KO and negative control infant splenic T cells (γδTCR^-^ FOXP3^-^ CD3^+^ cells) as determined by intracellular flow cytometry. Percentage of Helios-expressing cells in each group indicated on the top for both infant donors. **d**) msVIPER plots showing relative expression of genes from the Helios regulon (from ARACNe) in Helios KO relative to negative control cells by scRNA-seq. Genes positively regulated by Helios (“activated”) are in red and genes negatively regulated by Helios (“repressed”) are in blue. Top 6 genes from the Infant Tissue factor are labeled. **e**) Volcano plot showing FDR-adjusted *p*-value and log_2_ fold change in gene expression between Helios-KO and negative control CD8^+^ naïve/TCM from infant spleen from the TCR-stimulated condition. Data is averaged over both donor experiments for plotting and colored for genes with an FDR adjusted *p*-value < 0.05 and log_2_ fold change >1 (red) or <1 (blue) in both donors. **f**) Heatmap showing Z-scored average gene expression as log(counts per million+1) of up-regulated genes in Helios-KO versus negative control CD8^+^ naïve/TCM from the spleen in both infant donors.

We directly investigated the role of Helios in regulating its inferred targets by disrupting Helios expression in primary infant T cells using CRISPR-Cas9 gene editing. We isolated CD3^+^ T cells from the spleen of infant donors (ages 2 and 3 months old), transfected T cells with Cas-9 ribonucleoproteins targeting Helios without prior T cell stimulation, confirmed protein knockout (KO) by flow cytometry, and assessed the transcriptional profiles of Helios-KO infant T cells relative to controls at rest or following TCR-stimulation using scRNA-seq (**Fig. 6b**). CRISPR-Cas9 KO of Helios resulted in ∼60% reduction of Helios-expressing cells by flow cytometry for both infants (**Fig. 6c**). To validate the targets of Helios in our regulatory network, we assessed changes in gene expression of Helios-activated and Helios-repressed targets in Helios-KO CD8^+^ TEM relative to negative controls from the resting condition in the scRNA-seq data. We observed a marked inversion in expression of Helios’ gene targets in KO cells compared to controls, where Helios-activated targets were decreased and Helios-repressed targets were increased in both infants (**Fig. 6d**). These results experimentally confirm Helios’ ARACNe-inferred regulon, which is highly enriched in genes from the Infant Tissue factor that defines an infant-specific transcriptional state.

Following TCR-stimulation, we found only minor differences in gene expression for Helios-KO cells relative to controls in both CD4^+^ and CD8^+^ TEM (**Extended Data Fig. 3a,b**). However, Helios-KO in CD8^+^ naïve/TCM resulted in reduced expression of genes associated with IFN signaling (*MX1*, *GBP1*, *GBP4*, *GBP5*) and TFs regulating CD8^+^ memory T cell differentiation and function, including *ID2*^47^, *PLAC8*^48^, *HIC*^49^, *TOX2*^50^ (**Fig. 6e**). Conversely, we observed increased expression of an array of chemokines (*XCL1*, *XCL2*, *CCL3*, *CCL4*, *CCL4L2*, *CCL20*), pro-inflammatory cytokines (*IL2*, *CSF2*), cytotoxic mediators (*GZMB*), and co-stimulatory molecules (*TNFRSF9*, *TNFSF9*, *TNFRSF14*) (**Fig. 6f**). TCR-stimulated CD4^+^ naïve/TCM populations showed a similar pattern of increased expression for cytokines (IL2, CSF2, EBI3), chemokines (*XCL1*, *XCL2*) and co-stimulatory molecules and receptors (*TIGIT*, *KLRB1*, *TNFRSF4*, *TNFRSF9*, *TNFRSF18*, *TNFSF14*, *FCER1G*, *IL18R1*, *IL1R1*) in the Helios-KO condition relative to negative controls (**Extended Data Fig. 6c,d**). Together, our data demonstrate that Helios restricts effector functions of infant naïve/TCM cells after TCR-mediated activation, in addition to regulating a transcriptional program associated with T cell adaptation to tissues in infants.

## DISCUSSION

Our study provides in-depth transcriptional and functional analyses of tissue T cell responses and their underlying programming during a critical period of immune development in infancy relative to adulthood. We reveal unique transcriptional programs and multiple TFs expressed by infant memory T cells across tissues relative to adults. Infant memory T cells retain high levels of expression LEF1, TCF1, and KLF2, typically associated with stemness and quiescence among naïve T cells^51,52^. Also upregulated in infant memory T cells across tissues is the SRY-related HMG-box family TF SOX4, a critical transcriptional regulator that cooperates with TCF1 and LEF1 to facilitate T cell differentiation in the thymus^53^. Previous studies demonstrate that SOX4 regulates CD8^+^ memory T cell development^54^, antagonizes TH2 development in tandem with LEF1 in CD4^+^ T cells^55,56^, and facilitates stemness in cancer cells^57^, suggesting this TF network may play an important role in maintaining quiescence and differentiation potential among infant memory T cells. Furthermore, a recent study identified a small population of blood naïve T cells in healthy young adults expressing SOX4 and Helios (*IKZF2*) as recent thymic emigrants (RTEs)^58^. Several genes in RTE signature overlap with the top genes in the Infant Tissue signature (e.g., *TOX*, *SMC4*, *PDE7B, IKZF2*), raising the intriguing possibility that the infant-specific transcriptional state may arise from the generation of memory T cells from RTEs^59^.

Our findings demonstrate a globally reduced capacity for TCR-mediated activation among infant memory T cells relative to those in adults. This work expands on previous observations by our group and others showing that mucosal memory T cells from infants exhibit decreased production of inflammatory cytokines relative to older individuals^16,17,60^. These findings provide an intriguing contrast to our earlier work showing that naïve T cells from infants are more sensitive to TCR-stimulation and are biased towards differentiation into short-lived effector cells relative to adults^9,11^. Together, these data suggest that the infant immune system preferentially utilizes short-lived effector responses from naïve T cells to respond to infections, rather than effector memory responses in tissues.

We identify transcriptional programs strongly enriched in mucosal CD4^+^ and CD8^+^ TEM associated with tissue adaptation and residency. These findings extend and unify our previous work describing a shared tissue-associated signature in TEM from the bone marrow, lungs, and lymph nodes^19^ and tissue-specific adaptation signatures across barrier tissues^32^. Here, we elucidate a common tissue signature in TEM across multiple tissues in both infants and adults as well as a highly intestine-specific signature associated with gut homing, adhesion, and residency unique to adults. We identify KLF6 as a putative transcriptional regulator of both common and intestine-specific resident signatures using gene regulatory network inference. Accordingly, a recent study in mice also defines *Klf6* as a TF specific to the tissue-resident cell state using populational level RNA-seq and the Assay for Transposase-Accessible Chromatin using sequencing (ATAC-seq) data from a total of 10 murine studies^61^. Our trajectory analysis further suggests a role for KLF6 in the acquisition of a mature adult-like TRM phenotype over age, providing critical context to our previous work demonstrating staged maturation of TRM during infancy and childhood^16^.

Our study uncovers a distinct infant-specific transcriptional state in CD4^+^ and CD8^+^ TEM defined by expression of Helios (*IKZF2*), epigenetic modifiers *SMC4* and *CXXC5,* and regulators of tissue adhesion (*LAYN*, *CD9*). We provide direct experimental evidence of Helios’ role in driving the expression of genes in this transcriptional state using CRISPR-Cas9 knockdown, which also reveals Helios as a repressor of T cell effector function after TCR-stimulation. This finding is consistent with a previous report of human patients with Helios loss-of-function mutations that show enhanced IL-2 production by T cells from the peripheral blood^62^ or small populations of Helios-expressing T cells in the blood with reduced effector capacity^63^. Helios also promotes human fetal Treg differentiation and phenotype, where experimental knockdown of Helios enhances expression of pro-inflammatory genes in *ex-vivo* induced fetal Tregs^27^. Taken together, these data provide compelling evidence for tolerogenic state maintained by Helios in infant T cells.

This investigation reveals critical differences in transcriptional programming and activation capacity between infant and adult tissue T cells, with important implications for the generation and maintenance of protective immunity throughout the body. The expression of stem-like TFs in infant T cells may represent a developmental adaptation that facilitates rapid establishment of the tissue memory T cell niche. Conversely, the reduced activation capacity and effector function of infant memory T cells may serve to limit excessive inflammatory responses in tissues during this vulnerable period of development, but may also impair the ability to mount protective immune responses upon re-infection. Our work provides direct evidence for a cell-intrinsic mechanism regulating the infant-specific transcriptional state. This study lays the foundation for understanding the mechanisms governing infant T cells responses, which may aid identification of novel therapeutic targets to modulate T cell function in early life and promote long-lasting, protective adaptive immunity.

## METHODS

### Human organ donors and tissue acquisition

We obtained human organ tissue from deceased (brain-dead) organ donors directly at the time of acquisition for life-saving clinical transplantation through approved research protocols and materials transfer agreements with organ procurement organizations (OPOs) in the United States. Tissues from adult organ donors were obtained from an approved protocol with the OPO for the New York metropolitan area, LiveOnNY and included donors from our previous study^19^. Tissues from infant organ donors were obtained from LiveOnNY as well as through the Human Atlas for Neonatal Development (HANDEL) program based on the network for Pancreatic Organ Donors (nPOD). Consent for use of tissues for research was obtained by next of kin. All organ donors used in the study (**Supplementary Table 1**) were free of cancer, chronic disease, seronegative for hepatitis B, C and HIV and did not show evidence for active infection based on blood, urine, respiratory and radiological surveillance testing. The use of tissues from deceased organ donors does not qualify as human subjects research as confirmed by the Columbia University institutional review board.

### Tissue processing, T cell isolation and stimulation

Organ donor tissue samples were maintained in saline or University of Wisconsin solution on ice for transport to the laboratory and processing, typically within 2-24 hours of acquisition. Processing of infant and adult tissues to single cell suspensions was performed as previously described^16,19^. Briefly, blood was obtained by venipuncture, bone marrow was aspirated from the superior iliac crest, and mononuclear cells from both sites was obtained by density gradient centrifugation using Lymphocyte Separation Medium (Corning) or Ficoll-Paque Plus (Cytiva). Lymph nodes were isolated by dissection from the intestinal mesentery or the tracheobronchial tree of the lungs. Isolated lymph nodes, spleen and tonsil samples were placed in complete media composed of IMDM or RPMI 1640 (Gibco), 10% fetal bovine serum (GeminiBio), and 1% L-Glutamine:Pen:Strep Solution (GeminiBio), and mechanically dissociated with surgical scissors. Lung parenchymal tissue was dissected from the large airways, mechanically dissociated, and placed in digestion media composed of complete media, 1mg/ml Collagenase D (Millipore Sigma), 0.1 mg/ml DNase (Millipore Sigma) in a shaker at 37⁰C for 30 minutes. Intestinal tissues were dissected by their anatomical locations (jejunum, ileum, colon, Peyer’s patches), washed with sterile PBS (Corning) to remove luminal content, mechanically dissociated with scissors, and placed into digestion media in a shaker at 37⁰C for 30 minutes. Dissociated and/or digested cell suspensions of the lymph nodes, spleen, tonsils, lungs, and intestines were filtered using a 100 um filter (VWR) and centrifuged on a density gradient as above to remove debris and isolate mononuclear cells, followed by resuspension in complete media.

T cells from individual tissue single cell suspensions were enriched via magnetic negative selection (EasySep Human T cell Enrichment Kit; STEMCELL Technologies) followed by dead removal (Milteyni Biotec) resulting in +80-95% purity. We cultured 0.5-1 million T cell-enriched cells from each tissue for 16 hours at 37⁰C in complete medium, with or without TCR stimulation using ImmunoCult Human CD3/CD28 T Cell Activator (STEMCELL Technologies), after which dead cells were removed (as above) before single cell encapsulation.

### Single cell RNA-sequencing and data processing

T cell-enriched samples were loaded onto the Next GEM Chromium Controller using the Chromium Next GEM Single Cell 3’ Reagent kit v3.1 from 10x Genomics for single cell encapsulation and library construction as per manufacturer’s suggested protocols. Libraries were sequenced on an Illumina NovaSeq 6000, targeting ∼300M raw reads per sample (∼60,000 raw reads per cell).

scRNA-seq data were aligned and demultiplexed as described in Szabo et al^19^ using a publicly available pipeline (https://github.com/simslab/DropSeqPipeline8). Briefly, for each sample we trimmed read 2 to remove 3’-poly(A) tails (>7 A’s in length), discarded reads with fewer than 24 nucleotides remaining after trimming, and aligned the rest to GRCh38 (GENCODE v.24 annotation) using STAR v.2.5.0^64^. We assigned an address comprised of a cell-identifying barcode, unique molecular identifier (UMI) barcode, and gene identifier to each read with a unique, strand-specific exonic alignment. We followed the method in Griffiths et al.^65^ to filter the reads for index swapping and collapsed PCR duplicates using the UMIs after correcting sequencing errors in both the cell-identifying and UMI barcodes to generate an initial, unfiltered count matrix for each sample.

To identify cell-identifying barcodes that correspond to actual cells and to filter low-quality single-cell profiles, we used the methodology described in Zhao et al.^66^. Briefly, we used the EmptyDrops algorithm^67^ to remove cell-identifying barcodes that primarily contain ambient RNA. We then filtered the resulting count matrix to remove cell barcodes with high mitochondrial alignment rates (>1.96 standard deviations above the mean for a sample), high ratio of whole gene body to exonic alignment (>1.96 standard deviations above the mean for a sample), high average number of reads per transcript or transcripts per gene (>2.5 standard deviations above the mean for a sample), or cells where >40% of UMI bases are T or where the average number of T-bases per UMI is at least four.

### In silico T cell purification

While the EmptyDrops algorithm used above is adept at removing cell-identifying barcodes that correspond to ambient RNA, debris or molecular aggregates are more problematic^67^. We used a Gaussian mixture model (GMM)-based filter to further identify low-quality cells that could result from unfiltered index-swapping artifacts. We noticed that histograms of the number of transcripts per cell for some samples contained a lower mode, particularly for samples that were co-sequenced with samples external to this study (**Supplementary Fig. 3a**). To identify and isolate these modes, we modeled each sample’s log-scale transcript per cell distribution as a two-component GMM using *sklearn.mixture.GaussianMixture* (scikit-learn, version 0.21.3) and considered samples with a high-mode to low-mode ratio of at least 1.2 (on a log_2_-scale) as candidates for further filtering at the single-cell level. For these samples, we set a cutoff at three standard deviations above the mean for the high mode, using the square-root of the GMM’s estimate of the high-mode’s variance (*sklearn.mixture.GaussianMixture.covariance*), and removed cells below this cutoff. Differential expression analysis between the filtered and unfiltered cells showed that these low-coverage, filtered cells were likely contaminants and enriched in neural markers like *NNAT, BSN, NCAM2, NRXN1, GRIA1, GABRA1*, and *BDNF*.

With count matrices for high-quality cells in hand for each sample, we removed all non-T cells from the data, including contaminating cells, multiplets, and cells in which apparent T cell marker expression was likely an artifact of cross-talk^19^. We first performed unsupervised Louvain clustering using Phenograph^68^ with highly variable marker selection and k-nearest-neighbor graph construction as described in Levitin et al^69^. Highly variable marker selection was performed at the donor level and applied to each sample within a given donor. Next, we labeled each cluster as a putative T cell cluster (*t_1_*), contaminant cluster (*t_-1_*), or unknown cluster (*t_0_*) based on the average normalized expression of *CD3D*, *CD3E*, and *CD3G*. For each sample, we then examined the distribution of the number of reads per transcript (RPT), which estimates the number of PCR amplicons per transcript. PCR recombination can result in a multi-modal RPT distribution as described in Szabo et al^19^. To determine an RPT threshold for non-artifactual transcripts, we modeled this distribution as a two-component Geometric-Negative Binomial using the Scipy functions *scipy.stats.genom.pmf*, *scipy.stats.nbinom.pmf*, and *scipy.optimize.curve_fit* and took the intercept of the two components. We defined high-confidence T cells within each *t_1_* cluster as cells that expressed any of *CD3D*, *CD3E*, *CD3G*, *TRAC*, *TRBC1*, *TRBC2*, *TRDC*, *TRGC1*, and *TRGC2* with RPT greater than the sample-specific threshold. Next, we performed pairwise differential expression analysis (as described below) between the high-confidence cells in the *t_1_* cluster and the cells in the *t_-1_* cluster for each sample. We used this analysis to construct a contaminant gene list for each donor. We considered a gene to be contaminant-specific if it was at least 10-fold enriched in at least two *t_-1_* clusters across the donor with FDR<10^-5^ and if it was not enriched in more than one *t_1_* cluster with greater than 10% enrichment and FDR<10^-2^. For each sample, we computed the proportion of the contaminant gene list that each cell expresses above the sample-specific RPT, which we call *p_c_*. We then fit a truncated Gaussian distribution to the distribution of *p_c_* specifically for the high-confidence T cells from t_1_ clusters to establish a threshold proportion *p_t_* at three standard deviations above the mean of the fit. Finally, to call T cells, we identified cells in t_1_ clusters with any of *CD3D*, *CD3E*, *CD3G*, *TRAC*, *TRBC1*, *TRBC2*, *TRDC*, *TRGC1*, and *TRGC2* detected with RPT greater than the sample-specific threshold and *p_c_* < *p_t_* as T cells. We also identified cells in t_0_ clusters with any of *CD3D*, *CD3E*, *CD3G*, *TRAC*, *TRBC1*, *TRBC2*, *TRDC*, *TRGC1*, and *TRGC2* detected with RPT greater than the sample-specific threshold and *p_c_* < *p_t_* as T cells as long as the cluster’s first quartile for *p_c_* was also less than *p_t_*. **Supplementary Fig. 3b-g** contains a graphical depiction of each step in this procedure for a sample.

To validate the above procedure, we clustered all T cells that resulted with Phenograph and found no clusters with systematically lower expression of *CD3D*, *CD3E*, or *CD3G* (i.e., less than half of their mean expression). Furthermore, we clustered and performed differential expression analysis on all cells for each sample that were not called T cells to verify that most populations represented non-T cells and that all *CD3D*, *CD3E*, or *CD3G*-expression populations also expressed contaminants in common with non-T cells (i.e., possible multiplets). We did not find any clusters without clear contaminants.

### Consensus single cell Hierarchical Poisson Factorization

scHPF is a Bayesian algorithm for probabilistic factorization of scRNA-seq count matrices that produces highly interpretable factors or gene co-expression signatures^69^. Here, we applied the consensus implementation of scHPF, which generates and integrates many independent models of large scRNA-seq data sets, identifies recurrent factors from these models, and learns a final consensus model^18^. We used consensus scHPF to generate a single factor model for the entire T cell data set presented here including infant, adult, resting, and activated T cells from all tissue sites.

To construct the class-balanced dataset for scHPF, we randomly sampled 1,000 cells from each sample from tissues where we had at least one adult and one infant donor (blood, bone marrow, jejunum, jejunum lymph node, colon, lung lymph node, and spleen). To correct for coverage differences between samples, we downsampled the count matrices in this training set to the same mean number of transcripts per cell (2,086 transcripts/cell). We then filtered the training data to contain only protein coding genes detected in at least 1% of cells after downsampling. We additionally created an equivalently downsampled test set of cells that were not in the training set, with up to 200 cells for each experimental sample in the training data.

Due to scHPF’s highly multi-modal posterior on this complex dataset, we used the consensus approach described previously^18^, which allowed us to capture highly robust patterns of expression that consistently appear across scHPF training models with different random initializations, while still giving the model the freedom to approximate the parameter values the best explain the data. First, we ran scHPF with 10 random initializations for *k* =10 through 20 and selected three out of each set of 10 models with the lowest mean negative log likelihood on the training data for each value of *k*. Using Walktrap clustering, we identified 29 modules of similar factors that were observed in multiple models, and used their median gene weights to reinitialize scHPF and learn a refined, consensus model with 29 factors.

We evaluated the consensus-initialized model as compared to randomly initialized models with the same number of factors using the mean negative log-likelihood of the held-out test set. The consensus-initialized model had significantly better loss than any of the randomly initialized models (2.3953 with consensus-initialized vs 2.4006+/-0.0002 SEM for the 10 randomly initialized models). Thus, the consensus model achieved better log-likelihood for the held-out data, ensured that the factors were robust against random initializations, and effectively automated the selection of the number of factors *k*.

To project the full dataset onto the reference model obtained above, we downsampled all cells that were not included in training to have the same mean number of transcripts per cell as the training data. We then used the scHPF command *prep-like* to generate an appropriately filtered and formatted count matrix for these remaining cells and projected them into the consensus model using the scHPF command *project* with default parameters.

### T cell subset classification and analysis

We used scHPF’s embeddings in combination with gene expression values to annotate T cell subsets. CD4^+^ and CD8^+^ T cells can be difficult to distinguish based on transcriptional profiles alone due to transcript drop-out, particularly for CD4^+^ T cells, and because CD4 vs. CD8 status is highly correlated with effector status for the cells profiled here. This problem is further exacerbated in stimulated T cells where subset-specific markers are downregulated. Because scHPF breaks transcriptional profiles down into component expression programs, we can leverage its representations to distinguish between subsets, even for activated T cells.

We used a Naïve Bayes classifier on scHPF’s cell scores concatenated with expression values for several key markers: *CD4, CD8A, CD8B, CCL5, SELL, TRDC, TYROBP, CCR7, CTLA4*, and *FOXP3*. We first defined separate training sets for infant and adult resting T cells from within the scRNA-seq dataset based on co-expression of these markers for CD4^+^ naïve/TCM, CD8^+^ naïve/TCM, CD4^+^ TEM, CD8^+^ TEM, CD4^+^ Tregs, and γδ T cells, according to the scheme in **Supplementary Table 5**. Next, we generated a concatenated matrix comprised of the scHPF cell scores for the consensus model described above and the log-normalized expression of the above markers. Size factors for normalized counts were computed using the *computeSumFactors* function in *scran* as described by Lun et al^70^. We then standardized the resulting concatenated feature matrix using the *sklearn.preprocessing.StandardScaler* function (scikit-learn, version 0.23.2), trained a Naïve Bayes classifier separately for resting infant and adult T cells with *sklearn.naive_bayes.GaussianNB.fit*, and predicted the remaining resting infant and adult T cells separately using *sklearn.StandardScaler.transform*. We repeated this exact procedure for the infant and adult T cells separately, but this time including both resting and activated T cells and omitting the CD4^+^ Treg class. We could not classify CD4^+^ Tregs from the activated T cells in our dataset, because conventional T cells upregulate many canonical CD4^+^ Treg markers upon stimulation. Thus, this second round of classification allowed us to share information between the resting and activated T cells while obtaining an annotation for the activated T cells that excluded the CD4^+^ Treg class. Finally, for activated T cells, we used the annotation obtained from this second round of classification for downstream analysis. For resting T cells, we used this same annotation, but substituted the CD4^+^ Treg class for any cell classified as a CD4^+^ T cell in the second round of classification that was also classified as a CD4^+^ Treg in the first round.

We used several approaches to validating this classifier. First, we obtained excellent agreement between the expression patterns of canonical T cell subset markers and the classifier results as shown in **Fig. 1c**. Second, because the resting and activated T cells originate from matched samples, the number of resting and activated cells in each class should be roughly equal to each other for cells from the same sample. As shown in **Supplementary Fig. 4** the median absolute deviation between resting and activated cell frequencies for CD4^+^ and CD8^+^ T cells is 2.3 and 1.6%, respectively. Similarly, for naïve/TCM, TEM, and γδ T cells we obtain 3.4%, 3.1%, and 0.2%, respectively.

Our third approach was to estimate the accuracy of the classifier and therefore we applied it to a multi-tissue immune cell dataset from an organ donor from which we obtained CITE-seq data from blood, bone marrow, lung, lung lymph node, jejunum, and spleen^21,71^. We used the surface protein data from CITE-seq as an orthogonal ground truth to the corresponding RNA expression data from each cell to which we applied the Naïve Bayes classifier. First, we defined cells into high-confidence subsets based on the surface protein data. We defined these subsets as: CD4^+^ naïve (CD4^+^ CD8^-^ TCRγδ^-^ CD45RA^+^ CD45RO^-^ CCR7^+^ CD62L^+^ CD27^+^ CD25^-^), CD8^+^ naïve (CD4^-^ CD8^+^ TCRγδ^-^ CD45RA^+^ CD45RO^-^ CCR7^+^ CD62L^+^ CD27^+^), CD4^+^ TEM (CD4^+^ CD8^-^ TCRγδ^-^ CD45RA^-^ CD45RO^+^ CCR7^-^ CD62L^-^ CD27^-^ CD25^-^), CD8^+^ TEM (CD4^-^ CD8^+^ TCRγδ^-^ CD45RA^-^ CD45RO^+^ CCR7^-^ CD62L^-^ CD27^-^), CD4^+^ Treg (CD4^+^ CD8^-^ TCRγδ^-^ CD45RA^+^ CD45RO^-^ CCR7^+^ CD62L^+^ CD27^+^ CD127^-^ CD25^+^), and γδ T cells (CD4^-^ TCRγδ^+^) and ensured that these classes were mutually exclusive. To define positive and negative cell populations for each surface protein marker, we log-transformed the marker’s expression level (log_2_(counts per thousand +1)) and fit a two-component Gaussian mixture model to the transformed expression distribution (**Supplementary Fig. 1a**). For each fit, we computed defined three expression thresholds: 1.96 standard deviations below the mean of the higher mode Gaussian (*L_1_*), 1.96 standard deviations above the mean of lower mode (*L_2_*), and the local minimum of the Gaussian mixture fit between the means of the two components (*L_3_*). We then set our thresholds for marker-negative and positive subpopulations as *min(L_1_,L_2_,L_3_)* and *max(L_1_,L_2_,L_3_)*, respectively, to establish our ground truth T cell subsets. Finally, we trained a consensus scHPF model on the scRNA-seq component of the CITE-seq dataset, trained the Naïve Bayes classifier using the same procedure as described above based only on the scRNA-seq, and classified the high-confidence cells established using CITE-seq. Importantly, the Naïve Bayes classifier was blinded to the surface protein data used to define the high-confidence subsets and to the high-confidence subset annotations themselves. By comparing the Naïve Bayes classifier results to the high-confidence subsets established from surface protein expression, we found that the Naïve Bayes classifier was highly performant with favorable results for sensitivity, specificity, precision, and accuracy across T cell subsets (**Supplementary Fig. 1b**).

For downstream analysis, we interrogated T cells that were classified as CD4^+^ naïve/TCM, CD8^+^ naïve/TCM, CD4^+^ TEM, CD8^+^ TEM, CD4^+^ Tregs, and γδ T cells. We visualized the data using supervised dimensionality reduction and embedded the data into two dimensions by UMAP^20^, with T cell subsets as classes. Plotting UMAP embeddings and markers for each T cell subset was performed with *scanpy*.

### Differential gene expression between infant and adult T cells

We performed differential expression analysis across age groups for each conventional T cell subset (CD4^+^ naïve/TCM, CD8^+^ naïve/TCM, CD4^+^ TEM, CD8^+^ TEM, CD4^+^ Treg), using all tissue samples that met criteria of at least 100 cells per subset-donor-tissue combination. For CD4^+^ and CD8^+^ naïve/TCM, this included the blood, jejunum- and lung-associated lymph nodes, and spleen; for CD4^+^ and CD8^+^ TEM, this included the jejunum, lung and spleen; and for CD4^+^ Tregs, this included jejunum- and lung-associated lymph nodes as well as the spleen. For each tissue group within a subset, we performed pairwise differential expression using *scanpy v1.9.3* (*rank_genes_groups*; Wilcoxon with tie correction) between each infant donor versus every adult donor, using equalized cell counts (subsampled) and total counts (downsampled) for each group. We used the intersection of differentially expressed genes (FDR adjusted p-value < 0.05, log-fold change > 1) for every infant-adult comparison (i.e., must be differentially expressed in each infant donor compared to every adult donor) within each tissue to generate a list of differentially expressed genes across age in a given tissue. We next used the union of differentially expressed genes across all tissue comparisons within a subset to generate a final list of genes by T cell subset (**Supplementary Table 2**). UpSet plots were generated using the python package UpSetPlot *v0.8.0*.

### Activation trajectories by diffusion maps

We constructed activation trajectories for CD4^+^ and CD8^+^ TEM for mucosal tissues, where these subsets are highly enriched in TRM using diffusion component analysis as described previously^19^. Briefly, for CD4^+^ TEM and CD8^+^ TEM separately, we randomly sampled the same cell numbers from resting infant, resting adult, activated infant, and activated adult conditions from the mucosal sites lung, jejunum, ileum, and colon. Next, we computed a pairwise Euclidean distance matrix for the scHPF cell scores of the sampled mucosal CD4^+^ and CD8^+^ TEM, and used the DMAPS package (https://github.com/hsidky/dmaps) to embed the scHPF model into its first two diffusion components, which consistently separated the cells based on activation status.

### ARACNe and VIPER analysis

We used ARACNe-AP (https://github.com/califano-lab/ARACNe-AP) to infer gene regulatory networks from the scRNA-seq dataset^37,38^ using the metacell workflow described by Vlahos et al^72^. While ARACNe has been widely used for regulatory network inference from bulk RNA-seq data, the sparsity of scRNA-seq data requires the construction of pseudo-bulk profiles that average the expression profiles of multiple individual cells called metacells. To quantify cell-cell similarity for generating metacells, we used the cell score matrix from the consensus scHPF model described above into a Pearson correlation matrix from which we generated a k-nearest neighbors graph with k=50 to aggregate scRNA-seq profiles into metacell profiles by averaging over 50 similar cells as described previously^72^. Finally, we used ARACNe-AP to compute a transcription factor-target gene regulatory network consolidated from 200 rounds of bootstrapping using the metacell matrix and a list of 688 transcription factors that met the criteria for inclusion in our scHPF model (see above).

We used the gene regulatory network to associate transcription factors with scHPF factors. For this, we identified the subset of ARACNe-inferred targets that were activated by a given transcription factor using the *aracne2regulon* function in the VIPER package^46^. Then, we performed GSEA for each scHPF factor-transcription factor pair where the ranked gene list was obtained by ranking all genes by their scHPF gene score for a given scHPF factor and the gene sets were the set of activated targets for each transcription factor. This calculation yielded a normalized enrichment score reflecting the enrichment of a given transcription factor’s activated targets among the top-ranked genes in a given scHPF factor. We also used the gene regulatory network to infer transcription factor activities at the single-cell level using the VIPER algorithm (v1.26.0), a companion tool for calculating protein activity from ARACNe-inferred networks^46^. Specifically, we used the *viper* function with the z-scored, log-normalized scRNA-seq expression matrix (using *scran* as described above) to compute transcription factor activities with default parameters. Similarly, to assess differential VIPER activity between two conditions (e.g., for the Helios KO experiments described below), we generated a gene signature for the two conditions using the *rowTtest* function in VIPER, a null model using the *ttestNull* function in VIPER with 1,000 permutations, and the *msviper* function in VIPER with default parameters.

### Flow cytometry and intracellular staining

Single cell suspensions of tissue mononuclear cells washed with staining buffer comprised of PBS (Corning), 2% FBS (GeminiBio) and 2 mM EDTA (Gibco), incubated with Human TruStain FcX (BioLegend) for 10 minutes on ice. Cells were then stained with fluorescently labeled antibodies (**Supplementary Table 6**) for 30 minutes on ice and washed with staining buffer to remove unbound antibodies. We used the True-Nuclear Transcription Factor Buffer Set (BioLegend) for fixation, permeabilization and intracellular transcription factor antibody staining (**Supplementary Table 6**) according to manufacturer’s recommended protocols.

For stimulation assays, single cell suspensions of tissue mononuclear cells were cultured in complete media in 96-well U-bottom plates (Corning) at ∼2-5 million cells per well and stimulated with 50 ng/ml PMA (Sigma) and 1 ug/ml Ionomycin (Sigma) in the presence of GolgiStop and GolgiPlug (BD Biosciences) for 4 hours at 37⁰C. Cells were then washed with staining buffer and stained with surface antibodies as above. For fixation, permeabilization and intracellular cytokine antibody staining (**Supplementary Table 6**), we used the BD Cytofix/Cytoperm Fixation/Permeabilization Kit (BD Biosciences) as per manufacturer’s recommended protocols.

For all flow cytometry assays, we acquired cell fluorescence data using the Cytek Aurora spectral flow cytometer and analyzed data using FlowJo v10.10 (BD Life Sciences). For a gating strategy to identify T cell subsets refer to **Supplementary Fig. 5**.

### CRISPR-Cas9 deletion of Helios *(IKZF2)* in primary human tissue T cells

Mononuclear cell suspensions from infant spleens were obtained and T cell magnetic negative selection was performed as described above. For CRISPR-Cas9 deletion of Helios (*IKZF2*) we used a Cas9 RNP transfection approach^73^. Briefly, 3 Alt-R CRISPR-Cas9 crRNAs targeting Helios or negative controls (**Supplementary Table 6**) were individually complexed to Alt-R CRISPR-Cas9 tracrRNAs (IDT) in equimolar concentrations. Cas9 RNPs were prepared by combining crRNA:tracrRNA duplexes with TrueCut Cas9 Protein v2 (Thermo Fisher Scientific) at a molar ratio of 3:2. RNP nucleofection of T cells was performed using a Lonza 4D-Nucleofector X unit using the Lonza P3 Primary Cell 4D-Nucleofector X Kit with 3 ul of each RNP complex in 20 ul of nucleofection buffer and a pulse code of EH100. Nucleofected T cells were immediately placed in complete media with 5% human AB serum (Millipore Sigma) and cultured for 3 days at 37⁰C in an incubator for 3 days. Helios protein expression in Helios-KO T cells or negative controls was assessed after 3 days in culture by flow cytometry as described above. To assess the effects of Helios-KO on the T cell transcriptome, we first removed dead cells (Miltenyi Biotech) and either rested Helios-KO cells or negative controls overnight in complete media or stimulated cells with anti-CD3 and anti-CD28 and performed scRNA-seq as above.

We processed scRNA-seq data, identified T cells and categorized cells into T cell subsets using the Naïve Bayes classifier described above. We performed differential expression analysis between Helios-KO and negative control T cells in the activated condition, for each T cell subset with in each donor individually. We used the same method as above for identifying differentially expressed genes between KO and negative control cells (equalized cell numbers and counts; differential expression by Wilcoxon with tie correction). For visualizing differentially expressed genes (averaged FDR adjusted p-value < 0.05, averaged log_2_-fold change > 1) in volcano plots in **Fig. 6e**, we plotted averaged FDR-adjusted p-values and log-fold changes from both donors. Only genes that were differentially expressed between KO and negative control samples in both donors were plotted in **Fig. 6f**.

### Statistical Analysis

Descriptive analyses and statistical testing of flow cytometry data were performed using GraphPad Prism (v9.5.2) and comparisons between groups were made using statistical tests indicated in the figure legends. We considered comparisons as statistically significant for *p* < 0.05. For multivariate linear regression analysis between scHPF factor gene scores and relevant binary covariates (age, tissue type, subset, lineage, activation), we performed an ordinary least squares regression (statsmodels.OLS). Factor gene scores were z-scored for input and resulting regression coefficients were plotted for each factor and covariate.

## Data and Code Availability

Raw scRNA-seq data and metadata have been deposited in GEO (accession number GSE195844). Original source code with tutorials for scHPF, which is used to build individual models in consensus scHPF can be found at https://github.com/simslab/scHPF. Code for running consensus-scHPF along with helper scripts and instructions can be found at https://github.com/simslab/consensus_scHPF_wrapper.

## Acknowledgements

We extend our gratitude to the families of organ donors whose gift of donation made this work possible. We acknowledge the dedication and exceptional efforts of transplant coordinators and staff of LiveOnNY and the network for Pancreatic Organ Donors with Diabetes (nPOD) for tissue acquisition. This work was supported by US National Institutes of Health (NIH) grants to D.L.F. (AI106697, AI168634), D.L.F. and P.A.Si. (AI128949), and Helmsley Charitable Trust to D.L.F. and T.M.B. P.A.Sz. was supported by a Canadian Institutes of Health Research Fellowship and an AAI Intersect Fellowship. T.J.C. was supported by a K23 (AI141686) Award. J.J. was supported by the NIH Ruth L. Kirschstein Institutional National Research Service Award (5T32GM145440). D.P.C. was supported by the Columbia University Graduate Training Program in Microbiology and Immunology (T32AI106711).

Research reported in this publication was performed in the Columbia Center for Translational Immunology Flow Cytometry Core (supported by NIH S10OD030282) and the Sulzberger Columbia Genome Center and Single Cell Analysis core (supported by NIH P30CA013696). Experimental schematics in **Fig. 1a** and **Fig. 6b** were created with BioRender.com. Content is solely the responsibility of the authors and does not necessarily represent official views of the NIH.

## Author Contributions

Conceptualization: P.A.Sz, D.L.F., P.A.Si; Data Curation: P.A.Sz, H.M.L., T.J.C., P.A.Si; Formal Analysis: P.A.Sz, H.M.L., P.A.Si; Funding Acquisition: D.L.F., P.A.Si; Investigation: P.A.Sz, H.M.L, T.J.C., D.C., J.J., P.T., R.G., D.P.C., J.G., R.M., M.K; Methodology: P.A.Sz, H.M.L., P.A.Si; Resources: T.J.C., M.B., T.M.B., D.L.F., P.A.Si; Project Administration: D.L.F., P.A.Si; Supervision: D.L.F., P.A.Si; Visualization: P.A.Sz., D.P.C., P.A.Si; Writing – original draft: P.A.Sz, D.L.F., P.A.Si; Writing – review & editing: all authors.

## FIGURES

**Extended Data Figure 1:**
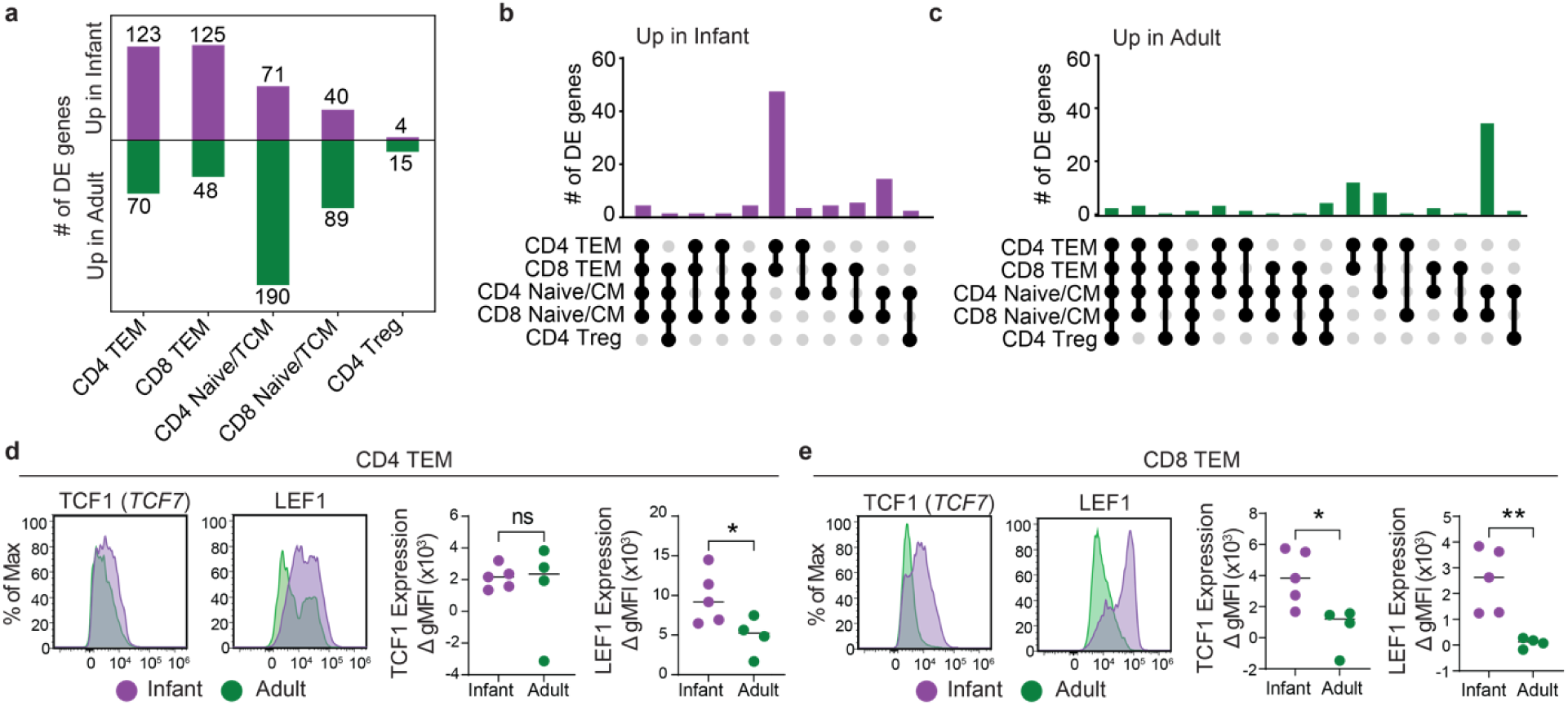
Differential gene expression analysis and TF staining by flow cytometry across infant and adult T cells. **a**) Bar plot showing number of differentially expressed genes upregulated in infants or adults by T cell subset across all tissues with adequate cell representation (**see Methods**). **b**) UpSet plot showing number of differentially expressed genes in infants relative to adults shared across at least two T cell subsets. **c**) UpSet plot similar to (**b**) but for differentially expressed genes upregulated in adults. **d**) Representative histograms showing TCF1 or LEF1 expression and quantification of geometric mean fluorescence intensity relative to isotype controls (ΔgMFI) in infant (n = 5) and adult (n = 4) CD4^+^ TEM from the spleen by flow cytometry. **e**) Representative histograms and quantification for TCF1 and LEF1 same as (**d**) but for CD8^+^ TEM. For panels (**d**) and (**e**), statistical comparisons between indicated groups made by Students’ *t*-test; “ns” denotes not significant, * *p* < 0.05, and ** *p* < 0.01.

**Extended Data Figure 2:**
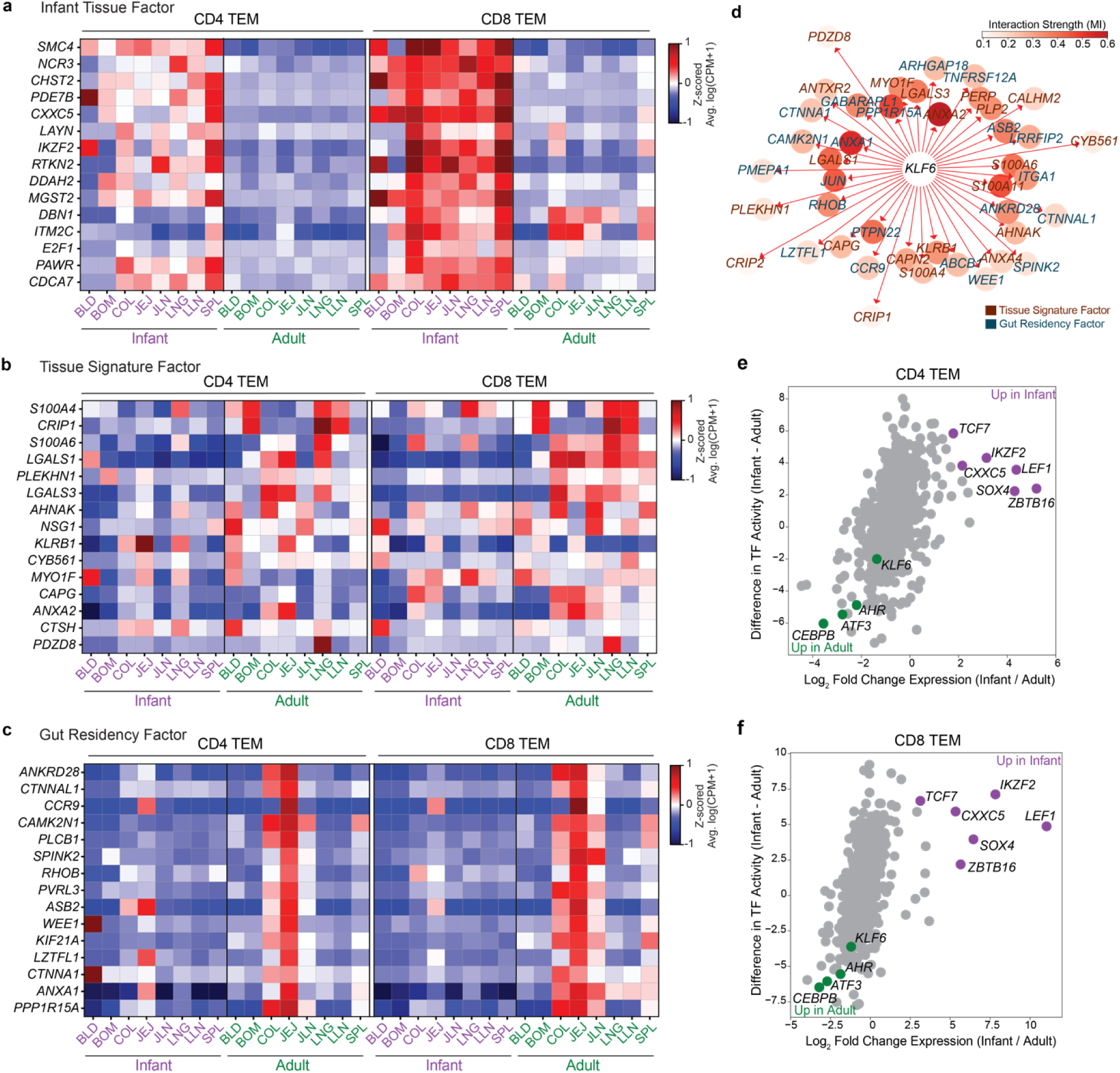
Expression of tissue-associated transcriptional programs in infants and adults. **a**) Heatmap showing Z-scored average gene expression as log(counts per million+1) of the top 15 genes in the Infant Tissue factor in CD4^+^ and CD8^+^ EM across tissues in infants and adults. **b, c**) Heatmap same as (**a**) but for the top genes in the Tissue Signature factor (**b**) and Gut Residency factor (**c**). **d**) TF network for KLF6 showing overlap of inferred targets within the top 50 genes in the Tissue Signature (*deep red*) and Gut Residency (*deep blue*) factors. Node color intensity and inverse distance in the network represents interaction strength (mutual information) between TF and target gene. **e, f**) Dot plots showing differences in TF activity (infant - adult) and log_2_ fold change in TF gene expression (infant / adult) in CD4^+^ (**e**) and CD8^+^ (**f**) TEM from the jejunum.

**Extended Data Figure 3:**
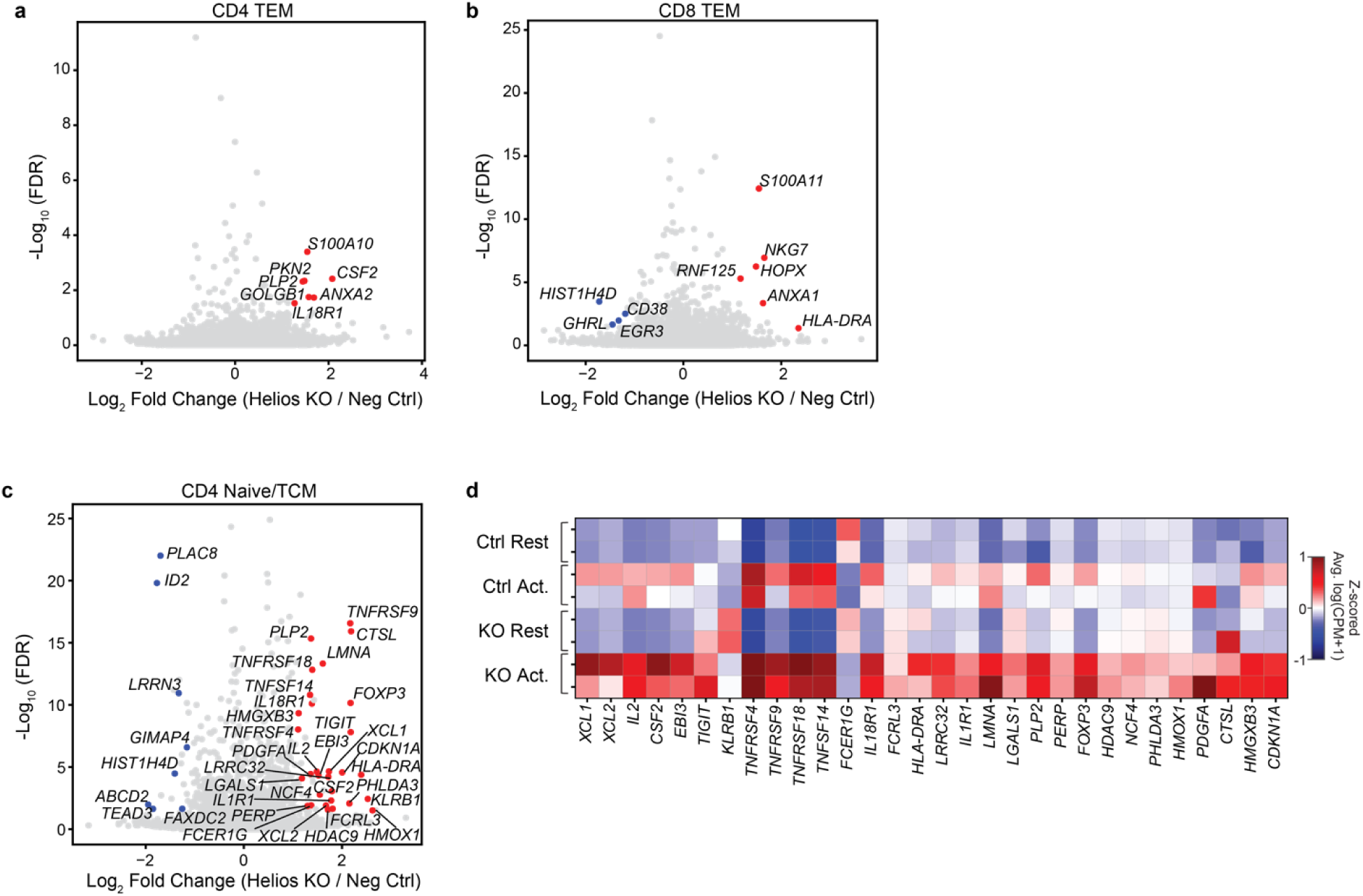
**Differential expression in Helios-KO versus negative control infant T cells. a,b**) Volcano plots showing FDR-adjusted *p*-value and log_2_ fold change in gene expression between Helios KO and negative control CD4^+^ TEM (**a**) and CD8^+^ TEM (**b**) from infant spleen in the activated condition. Data are averaged over both donor experiments for plotting and colored for genes with an FDR adjusted *p*-value < 0.05 and log_2_ fold change >1 (red) or <1 (blue) in both donors. **c**) Volcano plot as in (**a,b**) but for CD4^+^ naïve/TCM. **d**) Heatmap showing Z-scored average gene expression as log(counts per million+1) of up-regulated genes in Helios-KO versus negative control CD4^+^ naïve/TCM from the spleen in both infant donors.

