## Supplementary Figures and Tables for "Transcriptional control of T cell tissue adaptation and effector function in infants and adults"

**Supplementary Information for "Transcriptional control of T cell tissue adaption and effector function in infants and adults"**

**SUPPELMENTARY FIGURES**

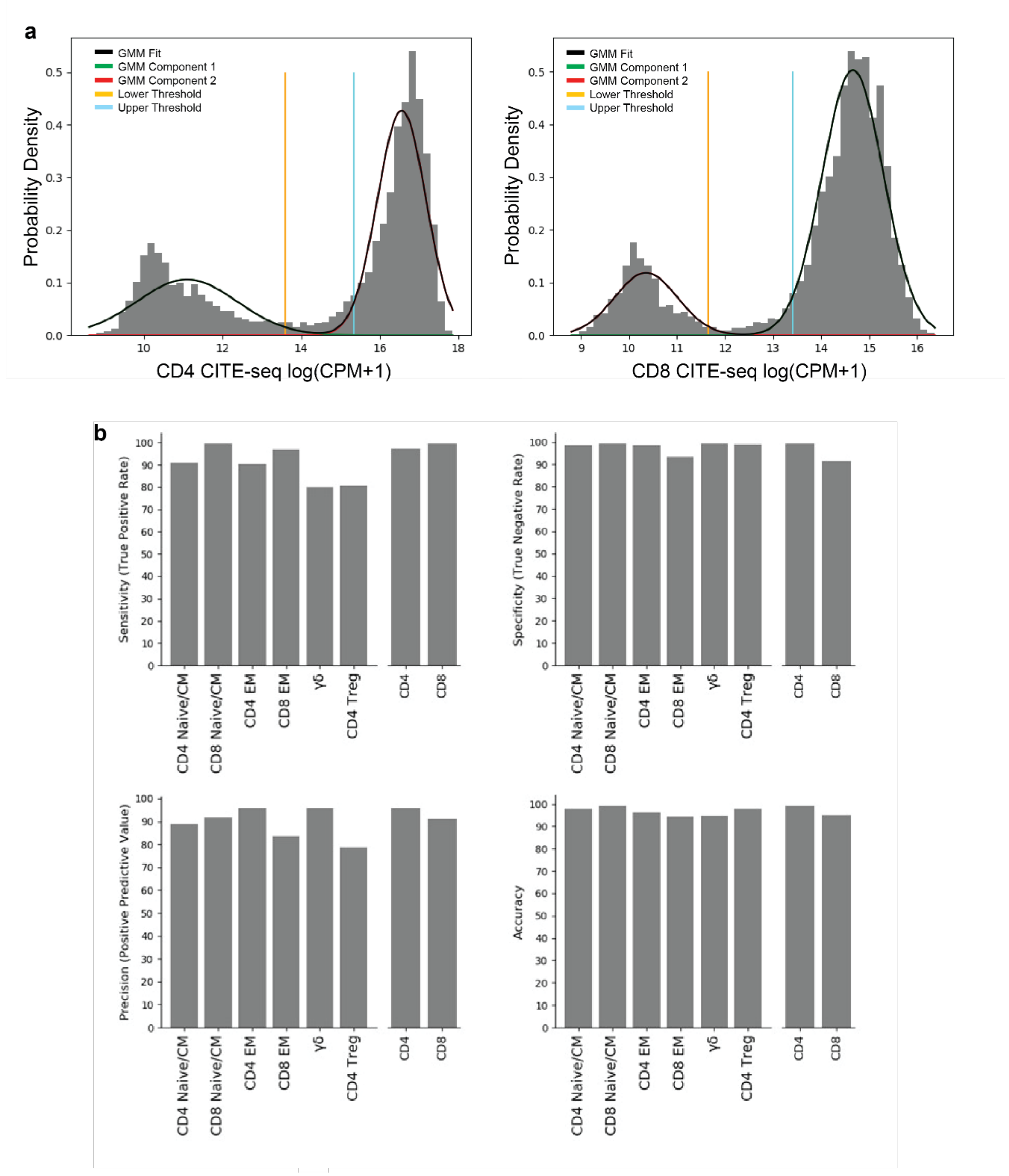

**Supplementary Fig. 1: Performance of Naïve Bayes T cell classifier on CITE-seq data from a human tissue atlas. a**) Expression distributions for CITE-seq data (obtained from a cross-tissue human immune atlas) with Gaussian mixture model fits and high-confidence thresholds for CD4 and CD8 surface markers. **b**) Sensitivity, specificity, precision, and accuracy for T cell subsets based on comparison of Naïve Bayes RNA-level classifier output and protein-level CITE-seq gating (used as a ground truth).

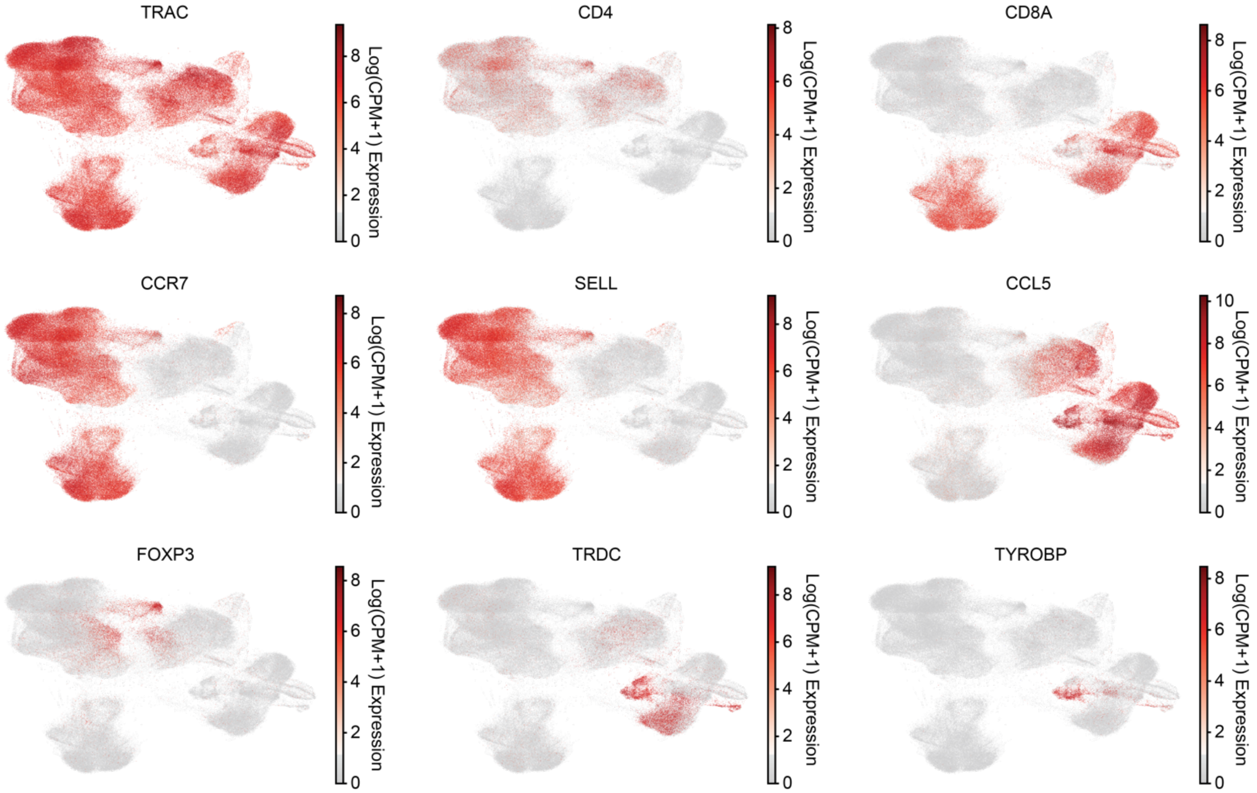

**Supplementary Fig. 2: Expression of T cell subset marker genes in the T cell activation map.** Indicated gene expression shown on a merged UMAP embedding of scRNA-seq profiles of T cells from infant and adult tissues.

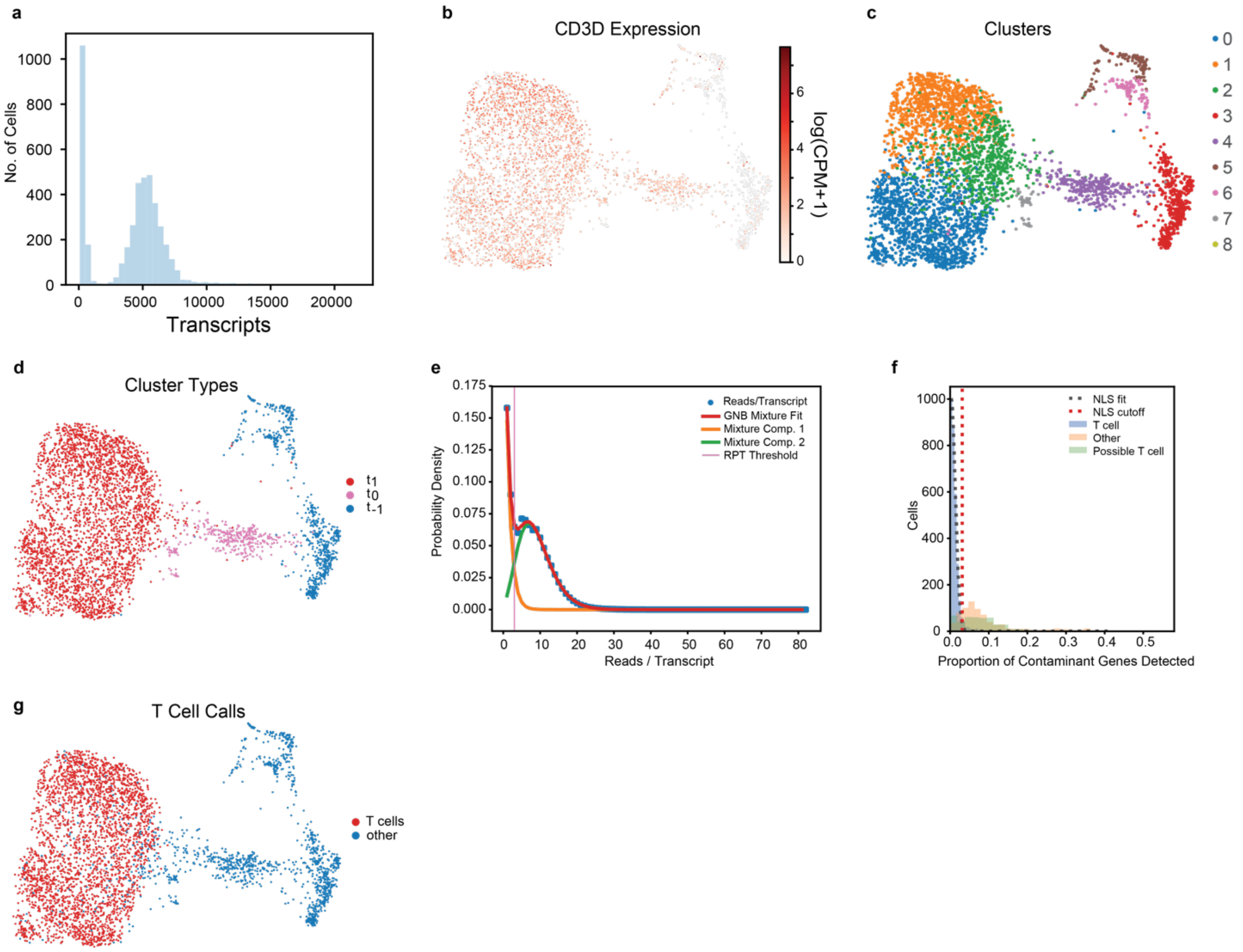

**Supplementary Fig. 3: T cell Calling. a**) Bimodal distribution of unique transcripts detected per cell for a representative sample. **b**) UMAP embedding of scRNA-seq data from a representative sample colored by expression of CD3D. **c**) Same as **b**) but colored by cluster identity. **d**) Same as **b**) but colored by cluster type (t1 = high-confidence T cells, t0 = high-confidence non-T cells, t-1 = possible T cells). **e**) Geometric-negative binomial mixture model fit to the distribution of reads per unique transcript with threshold; **f**) Non-linear least-squares (NLS) fit of truncated Gaussian distribution to the distribution in the proportion of the contaminant gene list detected per cell for high-confidence T cells with threshold (also shown are the same distributions for other cells and possible T cells); **g**) same as **b**) but colored by high-confidence T cell calls.

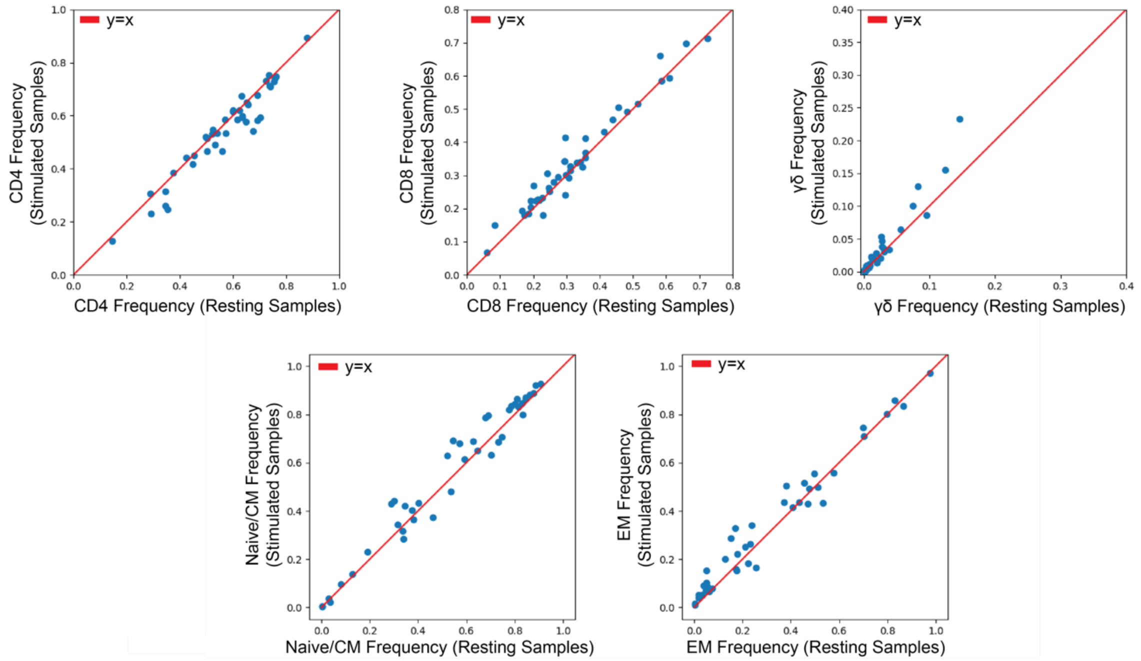

**Supplementary Fig. 4: Deviation between resting and activated T cell subset frequencies using the classifier.** Comparison of subset frequencies inferred by naive Bayes classifier for resting and stimulated tissue samples for CD4 T cells, CD8 T cells, γδ T cells, naive/central memory T cells, and effector memory T cells.

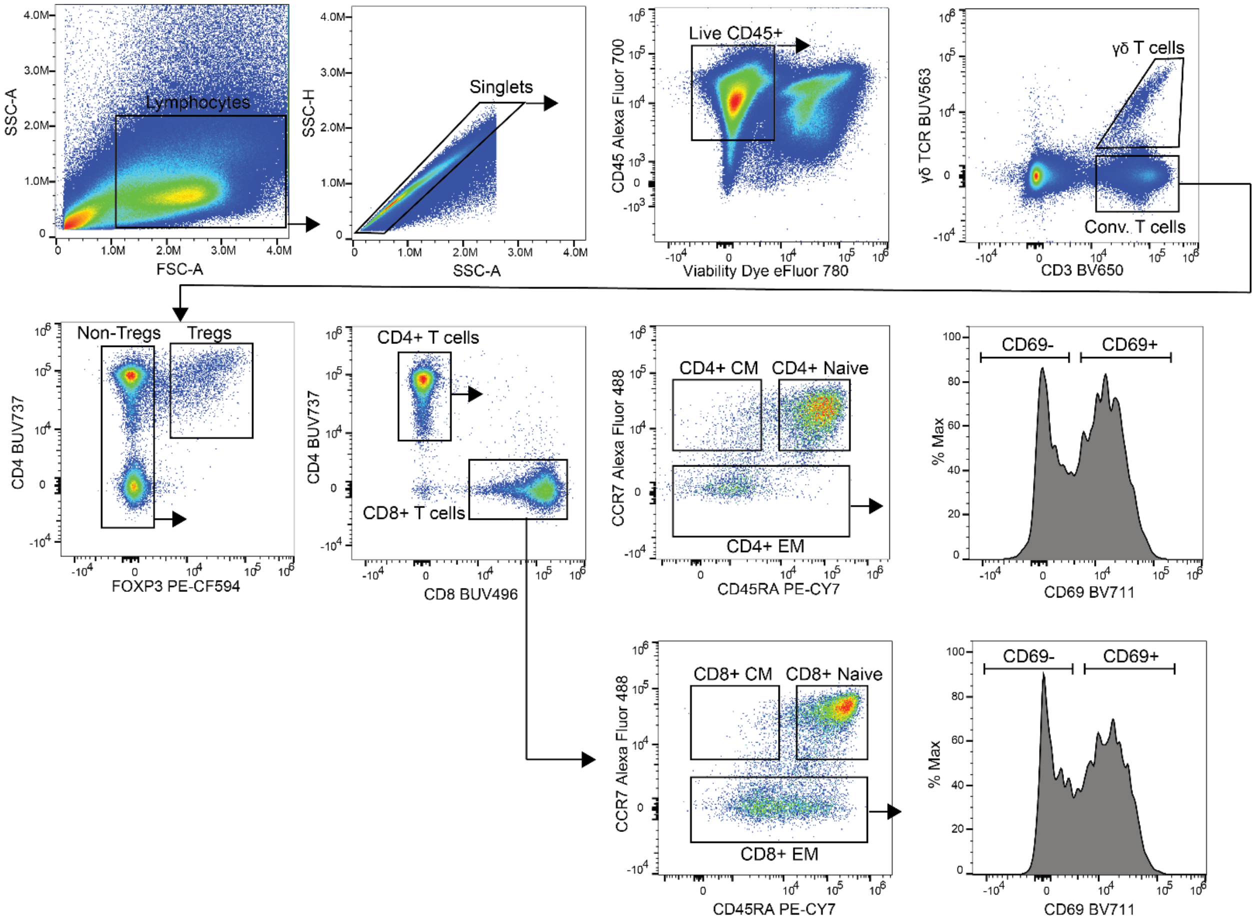

**Supplementary Fig. 5: Flow cytometry gating strategy for analysis of infant and adult tissue T cells.** Representative flow cytometry plots showing the gating strategy for T cell subsets in the spleen of an infant. Lymphocytes were gating based on forward scatter (FSC-A) and side scatter (SSC-A), singlets were gated on SSC-A and SSC-H. Live CD45$^+$ cells were gated on CD45 and viability dye exclusion. Conventional T cells were identified as CD3$^+$ and negative for γδ-TCR and Tregs were excluded based on intracellular expression of FOXP3. CD4$^+$ and CD8$^+$ T cells subsets were gated on CD4$^+$ or CD8$^+$ expression, respectively, and naïve cells were gated as CCD7$^+$ CD45RA$^+$, central memory cells were gated as CCR7$^+$ CD45RA$^-$, and effector memory cells were gated as CCR7$^-$ CD45RA$^{+/-}$. Expression of CD69 was gated on CD4$^+$ and CD8$^+$ TEM populations.

### SUPPLEMENTARY TABLES

#### Supplementary Table 1: Human organ donors and tissues for the study.

#### Donors used for scRNA-seq

| Donor ID | Age(yr) | Sex | COD | Blood | BM | Gut | LN | Lung | Spleen | Tonsil |
|---|---|---|---|---|---|---|---|---|---|---|
| Adult 1* (D363) | 63 | M | ICH | | X | | LLN | X | | |
| Adult 2* (D370) | 52 | M | Head Trauma | | X | | LLN | X | X | |
| Adult 3 (D457) | 40 | M | ICH | X | | JEJ, COL | JLN | | | |
| Adult 4 (D471) | 44 | M | Anoxia | X | | JEJ, COL | JLN | | X | |
| Infant 1 (D381) | 0.2 | M | Anoxia | X | X | | | | | X |
| Infant 2 (HDL049) | 0.2 | F | Anoxia | X | | JEJ, ILE, PEP | LLN, JLN | X | X | |
| Infant 3 (HDL055) | 0.25 | M | Head Trauma | | | JEJ, ILE, COL | LLN, JLN, CLN | X | | |
| Infant 4 (HDL082) | 0.75 | F | Head Trauma | | | JEJ, ILE, COL | LLN, JLN | X | X | |
| Healthy A* | 50s | M | N/A | X | | | | | | |
| Healthy B* | 50s | M | N/A | X | | | | | | |

*** from Szabo et al. Nat. Commun. 2019.**

**Donors used for Flow Cytometry and CRISPR-Cas9**

| Donor ID | Age(yr) | Sex | COD | Ext.Fig. 1 | Fig. 4 | Fig. 6A | Fig. 6C-F |
|---|---|---|---|---|---|---|---|
| Infant (LI004) | 0.25 | M | Anoxia | X | | X | X |
| Infant (HDL133) | 3 | M | Head Trauma | X | | | |
| Infant (HDL138) | 2 | F | Anoxia | X | | | |
| Infant (HDL148) | 2 | M | Head Trauma | X | X | | |
| Infant (HDL155) | 0.1 | M | Head Trauma | X | X | X | X |
| Adult (D555) | 64 | M | Anoxia | X | | X | |
| Adult (D559) | 64 | M | ICH | X | | | |
| Adult (D572) | 56 | M | Head Trauma | X | X | | |
| Adult (D587) | 58 | M | ICH | X | | | |
| Infant (HDL131) | 1.9 | M | Head Trauma | | X | X | |
| Adult (D570) | 73 | M | ICH | | X | | |
| Adult (D575) | 45 | F | ICH | | X | | |
| Infant (HDL133) | 3 | M | Head Trauma | | | X | |
| Infant (HDL110) | 0.4 | F | Anoxia | | | X | |
| Adult (D591) | 78 | M | Head Trauma | | | X | |
| Adult (D661) | 45 | M | Anoxia | | | X | |

**Abbreviations:** ICH, intracranial hemorrhage; JEJ, jejunum; ILE, ileum; COL, colon; PEP, Peyer's patches; LLN, lung-associated lymph node; JLN; jejunum-associated lymph node; CLN, colon-associated lymph node.

**Supplementary Table 2: List of differentially expressed genes across tissues by T cell subset for infants versus adults.** See attached file.

**Supplementary Table 3: Ranked list of top 100 genes for each scHPF factor.** See attached file.

**Supplementary Table 4: Transcription factor regulatory network for T cells generated by ARACNe.** See attached file.

**Supplementary Table 5: T cell subsets and markers for T cell classification in resting or activated conditions.**

| Resting Conditions | |
|---|---|
| **T cell Subset** | **Markers** |
| CD4 Treg | *CD4+/CD8-/FOXP3+/CTLA4+/CCL5-/TRDC-* |
| Adult CD4 Naïve/CM | *CD4+/CD8-/SELL+/CCL5-/TRDC-/FOXP3-/CTLA4-* |
| Adult CD8 Naïve/CM | *CD4-/CD8+/SELL+/CCL5-/TRDC-/FOXP3-/CTLA4-* |
| Adult CD4 EM | *CD4+/CD8-/SELL-/CCL5+/TRDC-/FOXP3-/CTLA4-/CCR7-* |
| Adult CD8 EM | *CD4-/CD8+/SELL-/CCL5+/TRDC-/FOXP3-/CTLA4-/CCR7-* |
| Adult γδ | *TRDC+/CCL5+/SELL-* |
| Infant CD4 Naïve/CM | *CD4+/CD8-/SELL+/CCL5-/TYROBP-/FOXP3-/CTLA4-* |
| Infant CD8 Naïve/CM | *CD4-/CD8+/SELL+/CCL5-/ TYROBP-/FOXP3-/CTLA4-* |
| Infant CD4 EM | *CD4+/CD8-/SELL-/CCL5+/TYROBP-/FOXP3-/CTLA4-/CCR7-* |
| Infant CD8 EM | *CD4-/CD8+/SELL-/CCL5+/ TYROBP-/FOXP3-/CTLA4-/CCR7-* |
| Infant γδ | *CCL5+/SELL-/CCR7-/TYROBP+/CD4-/FOXP3-/CTLA4-/CCR7-* |
| **Activated Conditions** | |
| **T cell Subset** | **Markers** |
| Adult CD4 Naïve/CM | *CD4+/CD8-/SELL+/CCL5-/TRDC-* |
| Adult CD8 Naïve/CM | *CD4-/CD8+/SELL+/CCL5-/TRDC-* |
| Adult CD4 EM | *CD4+/CD8-/SELL-/CCL5+/TRDC-/CCR7-* |
| Adult CD8 EM | *CD4-/CD8+/SELL-/CCL5+/TRDC-/CCR7-* |
| Adult γδ | *TRDC+/CCL5+/SELL-* |
| Infant CD4 Naïve/CM | *CD4+/CD8-/SELL+/CCL5-/TYROBP-* |
| Infant CD8 Naïve/CM | *CD4-/CD8+/SELL+/CCL5-/ TYROBP-* |
| Infant CD4 EM | *CD4+/CD8-/SELL-/CCL5+/TYROBP-/CCR7-* |
| Infant CD8 EM | *CD4-/CD8+/SELL-/CCL5+/ TYROBP-/CCR7-* |
| Infant γδ | *CCL5+/SELL-/CCR7-/TYROBP+/CD4-/CCR7-* |

**Supplementary Table 6: Antibodies and crRNAs used in the study.**

| ANTIBODIES | SOURCE | CLONE/IDENTIFIER |
|---|---|---|
| Anti-human CD45 Alexa Fluor 700 | BioLegend | 2D1; Cat: 368514 |
| Anti-human CD3 Brilliant Violet 650 | BioLegend | UCHT1; Cat: 300468 |
| Anti-human CD4 Brilliant Ultra Violet 737 | BD Biosciences | SK3: Cat: 612748 |
| Anti-human CD8 Brilliant Ultra Violet 496 | BD Biosciences | RPA-T8; Cat: 612942 |
| Anti-human CCR7 Alexa Fluor 488 | BioLegend | G043H7; Cat: 353206 |
| Anti-human CD45RA PE/Cyanine7 | BioLegend | HI100; Cat: 304126 |
| Anti-human CD69 Brilliant Violet 711 | BioLegend | FN50; Cat: 310944 |
| Anti-human γδTCR Brilliant Ultra Violet 563 | BD Biosciences | 11F2; Cat: 748534 |
| Anti-human FOXP3 PE-CF594 | BD Biosciences | 236A/E7; Cat: 563955 |
| Anti-human LEF1 (Primary Antibody) | Thermo Fisher Scientific | E.281.8; Cat MA5-14966 |
| Anti-human TCF-7/TCF-1 PE | BD Biosciences | S33-966: Cat: 564217 |
| Anti-human Helios PE | BioLegend | 22F6; Cat: 137216 |
| Anti-human CD45 APC-Fire 810 | BioLegend | HI30; Cat: 304076 |
| Anti-human TNF-α Brilliant Violet 785 | BioLegend | Mab11; Cat: 502948 |
| Anti-human IFN-γ Brilliant Violet 480 | BD Biosciences | B27; Cat: 566100 |
| Anti-human IL-2 PE/Dazzle 594 | BioLegend | MQ1-17H12: Cat: 500344 |
| Anti-human Granzyme B Alexa Fluor 700 | BD Biosciences | GB11; Cat: 560213 |
| Armenian Hamster Isotype Control PE | BioLegend | HTK888; Cat: 400907 |
| Rabbit IgG Isotype Control | Thermo Fisher Scientific | Cat: MA5-16384 |
| Mouse IgG1k Isotype Control PE | BD Biosciences | Cat: 554680 |
| Anti-rabbit IgG (H+L) Alexa Fluor 647 | Thermo Fisher Scientific | Cat: A21244 |
| **CRISPR-CAS9 crRNA ID** | **SOURCE** | **SEQUENCE/IDENTIFIER** |
| Alt-R CRISPR-Cas9 crRNA: Hs.Cas9.IKZF2.1.AA | IDT | CACCTCGCTGCTCTCAATTA |
| Alt-R CRISPR-Cas9 crRNA: Hs.Cas9.IKZF2.1.AB | IDT | CGAAAGGGAGCACTCCAATA |
| Alt-R CRISPR-Cas9 crRNA: Hs.Cas9.IKZF2.1.AL | IDT | CGAAAGGGAGCACTCCAATA |
| Alt-R CRISPR-Cas9 crRNA: Negative Control crRNA #1 | IDT | Cat: 1072544 |
| Alt-R CRISPR-Cas9 crRNA: Negative Control crRNA #2 | IDT | Cat: 1072545 |
| Alt-R CRISPR-Cas9 crRNA: Negative Control crRNA #3 | IDT | Cat: 1072546 |
